# Principles of chromosome organization for meiotic recombination

**DOI:** 10.1101/2024.03.06.583329

**Authors:** Mathilde Biot, Attila Toth, Christine Brun, Leon Guichard, Bernard de Massy, Corinne Grey

## Abstract

In meiotic cells, chromosomes are organized as chromatin loop arrays anchored to a protein axis. This organization is essential to regulate meiotic recombination, from DNA double-strand break (DSB) formation to their repair ^1^. In mammals, it is unknown how chromatin loops are organized along the genome and how proteins participating in DSB formation are tethered to the chromosome axes. Here, we identified three categories of axis-associated genomic sites: PRDM9 binding sites, where DSBs form ^2^, binding sites of the insulator protein CTCF, and H3K4me3-enriched sites. We demonstrated that PRDM9 promotes the recruitment of MEI4 and IHO1, two proteins essential for DSB formation ^3,4^. In turn, IHO1 anchors DSB sites to the axis components HORMAD1 and SYCP3. We discovered that IHO1, HORMAD1 and SYCP3 are associated at the DSB ends during DSB repair. Our results highlight how interactions of proteins with specific genomic elements shape the meiotic chromosome organization for recombination.

## INTRODUCTION

Meiotic recombination is initiated by DNA double-strand breaks (DSBs) that occur at PRDM9 binding sites in mice and humans ^5^. At the time of DSB formation, meiotic chromosomes are organized as an array of loops anchored to a chromosome axis composed of cohesins and structural proteins ^6^. The chromosome axis plays a role in mediating DSB repair regulation (e.g. choice of homologous chromosome *vs* sister and crossover *vs* non-crossover pathway) ^7^. It is, therefore, tempting to speculate that DSBs are formed in the context of the axis. In *S. cerevisiae*, it has been shown that sites prone to DSBs are located in loops, and are tethered to axes by multiple protein interactions ^8–11^. Furthermore, DSB formation requires evolutionarily conserved proteins located on chromosome axes ^12,13^. However, it is unknown how meiotic DSB sites and activity are coordinated with the chromosome spatial organization in mammals.

In mice, chromosome axes assemble during leptonema through interactions between cohesins and meiotic-specific proteins, such as SYCP2, SYCP3 and HORMAD1 ^14,15^. Interactions between PRDM9 and several axis proteins have been detected ^16^. Moreover, the essential DSB proteins REC114, MEI4 and IHO1 (RMI complex) are visible as axis-associated foci at leptonema, potentially restricting DSB formation near axes ^4,17,18^. These proteins form a complex ^19^, can form DNA-driven condensates *in vitro* ^20–22^, and are thought to promote DSB activity through direct interaction with the catalytic complex composed of SPO11 and TOPOVIBL ^23^. However, critical issues are: what determines the genomic and spatial localization of the RMI complex; how PRDM9 sites are recruited to axes and to the RMI complex; and whether there is a specific genomic organization of the loop-axis array. Hi-C maps of meiotic prophase spermatocytes have detected contacts, supporting the loop-axis organization; however, there are only few specific contact points along the genome, possibly due to the high contact complexity and a stochastic distribution among meiotic cells, thus leaving these questions unanswered ^24–29^.

Therefore, we analyzed by chromatin immunoprecipitation followed by next-generation sequencing (ChIP-seq) the genomic localization of two axis proteins (SYCP3 and HORMAD1) and two RMI proteins (MEI4 and IHO1) in mouse spermatocytes at the time of DSB formation. We also studied the molecular mechanism of their association with chromatin using different mouse mutants. This approach allowed us to: (i) discover that PRDM9 enables the recruitment of MEI4, IHO1 (in a MEI4-dependent manner) and the axis proteins (in an IHO1-dependent manner) to its sites; (ii) provide direct molecular evidence that axis proteins interact with DSB ends; and (iii) identify CTCF and H3K4me3 enriched-sites as preferentially axis-associated. These features uncover fundamental properties of the axis-loop organization of meiotic chromosomes, and provide a framework for understanding the coordination between the initiation of meiotic recombination and chromosome organization.

## RESULTS

### PRDM9 directs MEI4 to DSB sites

Cytological analysis of the localization of the RMI proteins REC114, MEI4 and IHO1 in mice showed several hundred axis-associated foci at leptonema, the stage of DSB formation. The number of foci decreases in parallel with DSB formation and synapsis. In the absence of DSB activity, MEI4 foci form and remain stably axis-associated ^3,4,18^. We first examined MEI4 genomic localization by ChIP-seq of purified chromatin from testes with synchronized spermatogenesis of juvenile C57BL/6 mice, called wild type (WT) hereafter. Meiosis synchronization allowed obtaining leptotene/zygotene spermatocytes (Figures S1A, B, Table S1). We normalized the ChIP signal obtained in WT samples using data from parallel experiments (same synchronization and ChIP-seq protocols) performed in a *Mei4KO* mouse strain. For peak calling, we used the Irreproducible Discovery Rate method and retrieved only peaks present in both replicates. We used this experimental strategy for all ChIP-seq experiments unless otherwise stated (see Star Methods, Table S1).

MEI4 showed a specific enrichment at multiple genome locations (Figure 1A). First, we asked whether the retrieved peaks (n=1566; Table S2) overlapped with DSB hotspots. In mice, DSB hotspots have been mapped by assessing the DMC1 recombinase enrichment by single strand DNA ChIP-seq (DMC1-SSDS) ^30^. The position of maximum DMC1-SSDS intensity corresponds to the hotspot center that also coincides with the maximum signal intensity of PRDM9 binding ^31^. The DMC1-SSDS data used here were obtained in a C57BL/6 mouse strain, carrying the same *Prdm9* allele as our samples (*Prdm9^Dom^*^2^). Differently from what reported in *S. cerevisiae* where MEI4 is mostly enriched at non-hotspot sites ^10^, 96.5% of the 1566 mouse MEI4 peaks localized at hotspots (Figure 1B, Table S3). We also detected a strong MEI4 enrichment within the chromosome X, 4, 9 and 13 subtelomeric regions. This correlates with previous cytological analyses showing high MEI4 accumulation in these regions that contain a high copy number of the mo-2 minisatellite ^32^ (Figures S1C-G, see Star Methods). At hotspots, the maximum intensity of the average MEI4 signal coincided with the hotspot center (Figure 1C), and MEI4 enrichment intensity was positively correlated with that of PRDM9, H3K4me3 and DMC1-SSDS (Figure 1D). We then tested the MEI4 signal in a *Spo11KO* strain where no DSB is formed ^33,34^. We still detected MEI4 binding at the hotspot center (Figure 1C, Figure S1H). In fact, MEI4 enrichment was stronger in *Spo11KO* than in WT spermatocytes, and both peak number (3817 vs 1566) and signal intensity were increased (Figure 1E, Figure S1I, Table S2, S4). For this comparison between genotypes and for subsequent analyses, we measured the median signal intensity over the 2000 strongest hotspots and the Cohen’s D (a standardized measure of the mean difference between samples) (see Star Methods and Table S4). Interestingly, the difference of the MEI4 ChIP signal between *Spo11KO* and WT mice was consistent with the cytological data: in WT spermatocytes, the number of MEI4 foci starts to decrease after leptonema, as MEI4 is displaced upon DSB formation ^17^. Consistently, in *Spo11KO* spermatocytes the number of MEI4 foci was higher than in WT spermatocytes (Figure S1J, upper panel). Moreover, MEI4 foci appeared brighter. Indeed, the median focus intensity of axis-bound MEI4 was increased by ∼4.6-fold at mid-leptonema in *Spo11KO* vs WT spermatocytes, indicating MEI4 signal accumulation at a given site (Figure S1J, lower panel). The observed increase in focus intensity at the cytological level is compatible with the read enrichment increase at hotspots, suggesting that MEI4 detected by ChIP, corresponds to the MEI4 foci seen by cytology.

**Figure 1.**
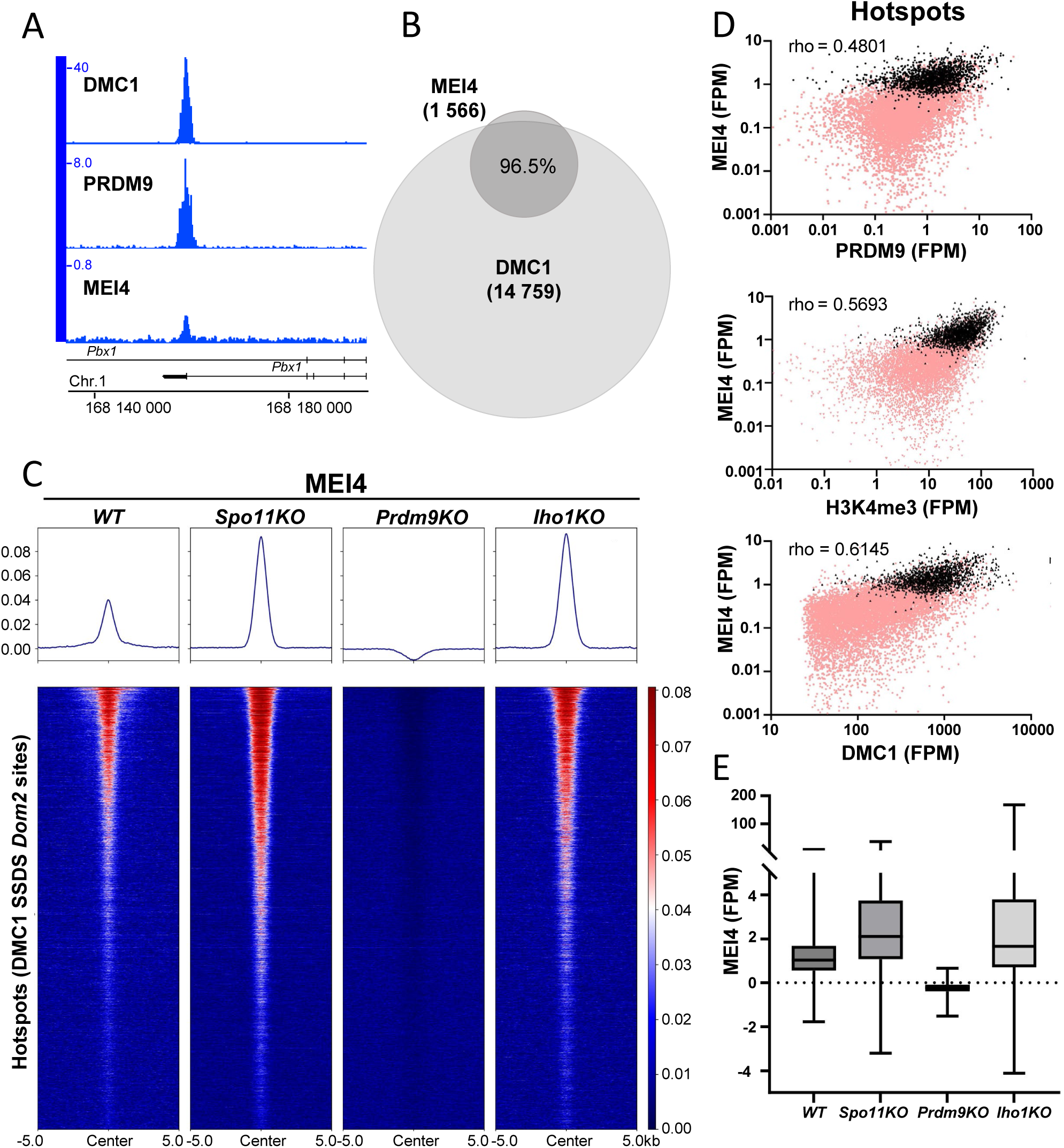
MEI4 is recruited to DSB sites in a PRDM9 methyltransferase dependent and IHO1 independent manner. **(A)** ChIP-seq read distribution for MEI4, DMC1-SSDS and PRDM9 ^31^ at a representative *Dom2* hotspot in wild type (WT) spermatocytes. **(B)** Venn diagram showing the overlap of MEI4 ChIP-seq peaks (dark gray) and DMC1-SSDS peaks^31^ (light gray) in WT (*B6)* mice. **(C)** MEI4 ChIP-seq signal in *WT*, *Spo11KO, Prdm9KO* and *Iho1KO* spermatocytes at all *Dom2* hotspots. DMC1-SSDS signal intensity was from ^31^. **(D)** MEI4 signal (FPM) compared with the PRDM9 signal intensity (FPM), H3K4me3 signal intensity ^86^, and DMC1-SSDS peak intensity ^31^ at all *Dom2* DSB hotspots in WT spermatocytes ^31^. Black and pink dots highlight hotspots that overlap and do not overlap with MEI4 peaks, respectively. rho: Spearman correlation coefficient. **(E)** MEI4 ChIP-seq signal intensity at *Dom2* hotspots ^31^ in *wt*, *Spo11KO*, *Prdm9KO* and *Iho1KO*spermatocytes. The signal was measured at the 2000 strongest hotspots.

Next, we monitored MEI4 enrichment at hotspots in *Prdm9KO* and showed a PRDM9-dependent recruitment (Figure 1C). However, in *Prdm9KO*, MEI4 forms robust foci, at similar levels as in WT spermatocytes ^35^. This paradox could be explained by the fact that in *Prdm9KO* mice, DSB activity does not take place at PRDM9 sites, but at sites corresponding mainly to promoters and enhancers, called default sites ^36^. However, closer inspection of the MEI4 signal on the genome browser and by peak calling revealed almost no signal, even at default sites, and peak calling detected only 80 peaks (Figure S1K, Table S2). This puzzling finding led us to test whether DSB formation in *Prdm9KO* mice was MEI4-independent. We generated *Mei4KO Prdm9KO* mice and examined DSB formation by immunostaining in male meiocytes. We observed low γH2AX signal levels and no RPA foci (two DSB repair activity markers), indicating the absence of DSB formation in *Mei4KO Prdm9KO* mice (Figure S2). Thus, in the absence of PRDM9, MEI4 is required for DSB formation, forms foci on chromosome axes, but is not detected by ChIP-seq in our conditions (S3A, S1K). The lack of MEI4 detection by ChIP at default sites in *Prdm9KO* could certainly be a sensitivity issue. At least two factors could contribute to a distinct signal detection between WT and *Prdm9KO*: i) the mode of MEI4 recruitment, with a potentially more distant chromatin localization in *Prdm9KO*, and ii) the number of potential genomic DSB sites is greater in *Prdm9KO*. To determine whether PRDM9 methyltransferase activity was required for MEI4 binding to hotspots, we analyzed MEI4 localization in the B6-Tg(YF) mouse strain. This strain produces two PRDM9 variants with distinct DNA binding specificities: PRDM9^Dom2^ with WT methyltransferase activity, and PRDM9^Cst^-YF with defective methyltransferase activity due to a point mutation (Y357F) in the PR/SET domain ^37,38^. We previously showed that PRDM9^Cst^-YF binds to the binding sites of PRDM9^Cst^, but does not catalyze the methylation of the surrounding histones ^37^. As expected, PRDM9 bound to both target sites (*Prdm9^Dom^*^2^ and *Prdm9^Cs^*^t^) (Figure S3B, left panels). However, MEI4 was only detected at sites bound by the catalytically active PRDM9^Dom2^ isoform (Figure S3B, right panels). The very weak remaining signal observed at the sites bound by PRDM9^Cst^-YF could be explained by a residual methyltransferase activity in this mutant, as previously suggested ^37^, or to some MEI4 binding to the catalytically inactive PRDM9. The positive control for MEI4 binding at these sites was the mouse RJ2 strain that expresses only PRDM9^Cst^ (Figure S3C). These results show that PRDM9 catalytic activity and likely the associated histone modifications are essential for MEI4 binding to hotspots.

In *S. cerevisiae,* Mer2 (the IHO1 ortholog) is required for DSB formation and is essential for Mei4 (and Rec114) recruitment to axes and DSB sites ^10^. Surprisingly, mouse MEI4 enrichment at DSB hotspots was not affected by IHO1 absence (Figure 1C). The signal was actually stronger than in WT (Figure 1E), possibly due to MEI4 persistence at its genomic sites due to the absence of DSB activity in this mutant, as observed in *Spo11KO* mice. A previous cytological study showed that in *Iho1KO* spermatocytes, the number of MEI4 foci is reduced by 11-fold compared to WT ^4^. In the light of our findings, we re-evaluated this quantification. We also found that the number of on-axis MEI4 foci was reduced in *Iho1KO* compared with WT and that conversely, the number of off-axis MEI4 foci and their intensity were increased (Figure S3D). Therefore, we propose that in the absence of IHO1, the main location of the MEI4-hotspot association detected by ChIP is not on chromosome axis. These observations imply that the axis-association of MEI4-bound hotspots depends on IHO1, and that MEI4 binding to hotspots is not sufficient for DSB formation.

### HORMAD1 and SYCP3 are localized at hotspots and on resected DSB ends

To gain insight into the axis organization, we determined the genomic localization of the two axis proteins SYCP3 and HORMAD1. SYCP3 binds to DNA, forms filaments by self-assembly ^39^ and forms a complex with SYCP2, which interacts with the two HORMAD paralogs 1 and 2 ^40^. SYCP3 and HORMAD1 are visible at pre-leptonema as small stretches or distinct foci, and partially colocalize with the kleisin subunits REC8 and RAD21L before the axial core structure is fully formed ^15,41–44^. SYCP3 remains axis-associated throughout prophase I, whereas HORMAD1 is progressively displaced from axes engaged in synapsis, starting at zygonema. SYCP3 is not required for DSB formation, but is important for efficient DSB repair and synaptonemal complex formation ^45,46^. In contrast, in *Hormad1KO* spermatocytes, DSB level is reduced by 4-fold, resulting in homologous synapsis defects ^47^.

We found that HORMAD1 and SYCP3 were enriched at specific genomic locations (Figure 2A). We detected 1715 (HORMAD1) and 1911 (SYCP3) peaks, among which 41% and 60% (706 and 1144 peaks), respectively, overlapped with DSB sites (Figure 2B, Table S2 and Figure S4). Like for MEI4, their intensity was positively correlated with that of DSB activity monitored by DMC1-SSDS (Figure 2C, Figure S4A). The HORMAD1 and SYCP3 enrichment at hotspots covered intervals up to 5 kb, with an average enrichment that unexpectedly showed a triple peak pattern (one central peak and two flanking peaks), with a maximum enrichment at about 1.5 kb from the hotspot center (Figure 2C). In the mouse, DSB ends are resected (∼1 kb, on average) at both sides ^48,49^. Direct comparison of the distribution of HORMAD1-associated hotspots and SYCP3 signals with the distribution of resection end tracts revealed that both axis proteins were positioned immediately adjacent and distal to the end-resection peak. This is compatible with an enrichment at or near the dsDNA-ssDNA transition (Figure 2D, Figure S4B) and provides direct evidence that after DSB formation, one or both sequences flanking resected DNA ends are axis-associated. We observed this pattern also for hotspots on the X chromosome where, unlike on autosomes, DSBs are thought to be mainly repaired by recombination with the sister-chromatid (Figure S4C). As the analyzed cell population contained mostly spermatocytes at zygonema (Table S1), we hypothesized that the triple peak signal at hotspots originated from two cell types: cells where DSB had not occurred yet (the central peak), and cells where DSB formation and resection had occurred (the flanking peaks). To test this hypothesis, we monitored HORMAD1 and SYCP3 hotspot enrichment in the absence of DSB activity (i.e. in *Spo11KO*, *Mei4KO* and *Iho1KO* mice).

**Figure 2.**
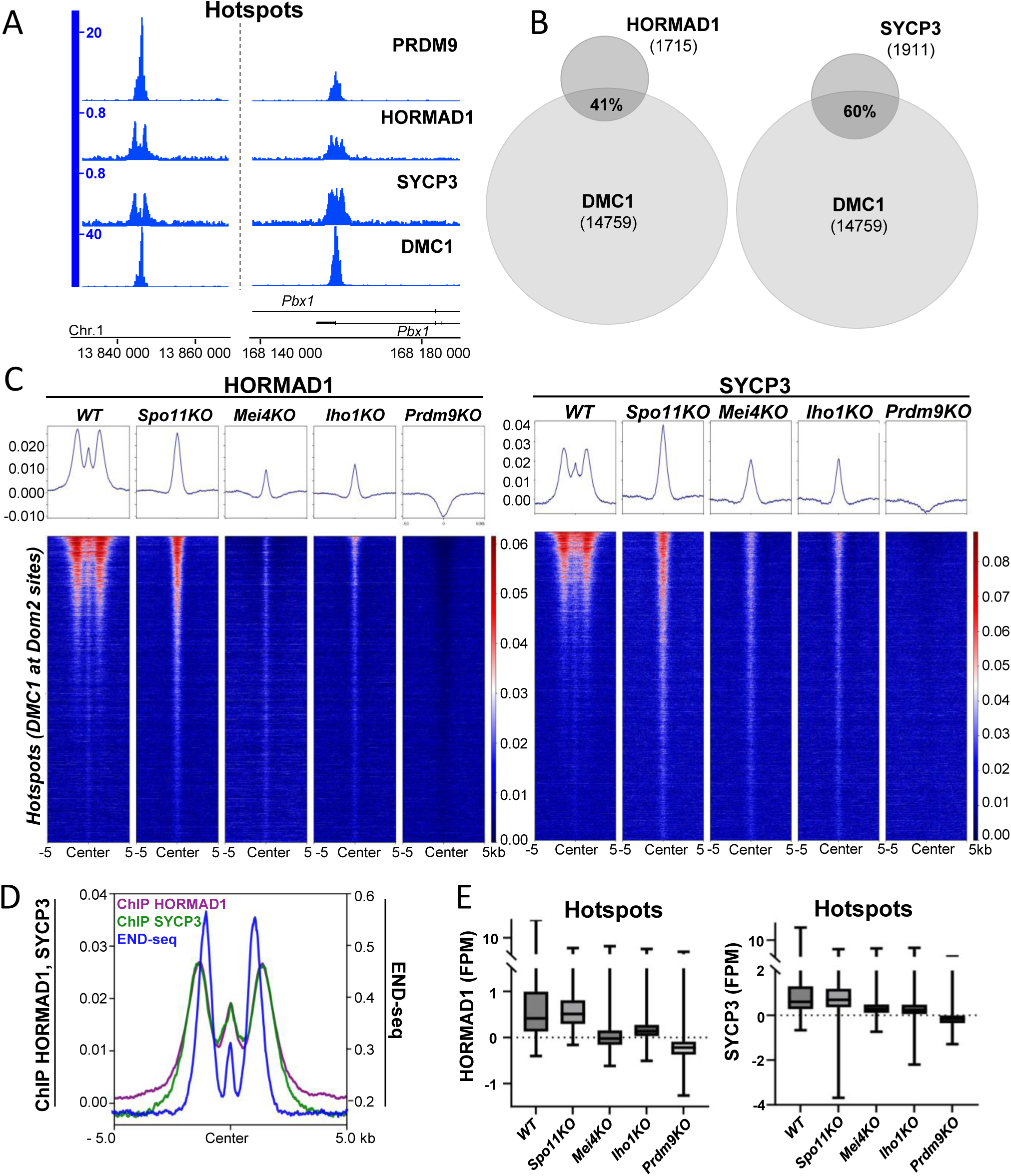
Two modes of interaction with hotspots for HORMAD1 and SYCP3. **(A)** ChIP-seq read distribution of HORMAD1 and SYCP3, DMC1-SSDS and PRDM9 at two representative *Dom2* hotspots in wild type (WT) spermatocytes. **(B)** Venn diagrams showing the overlap of HORMAD1 and SYCP3 ChIP-seq peaks (dark gray) and DMC1-SSDS peaks (light gray) ^31^ in WT mice. **(C)** HORMAD1 and SYCP3 ChIP-seq signal in *WT*, *Spo11KO, Mei4KO*, *Iho1KO* and *Prdm9KO* spermatocytes at all *Dom2* hotspots. DMC1-SSDS signal intensity was from ^31^. **(D)** Averaged enrichment of HORMAD1 (blue), SYCP3 (green) (ChIP-seq) and resection track ends, assessed by END-seq (red) ^48^, at *Dom2* hotspots ^31^. **(E)** HORMAD1 and SYCP3 ChIP-seq signal intensity at *Dom2* hotspots ^31^ in *Wt*, *Spo11KO*, *Mei4KO, Iho1KO* and *Prdm9KO* spermatocytes. The signal was measured within ±700bp from the center of the 2,000 strongest hotspots ^31^.

In *Spo11KO* mice, HORMAD1 and SYCP3 were present at hotspots (698 and 598 peaks representing 24% and 25% of all peaks, respectively), with an average enrichment as a single central peak (Figure 2C, Tables S2 and S3). The percentage of peaks within hotspots was lower in *Spo11KO* than in WT spermatocytes (41% and 60%, respectively) due to the wider and stronger signal associated with DSB end-resection in WT samples, as described above (Table S3, Figure S4D). For both HORMAD1 and SYCP3, the median intensity of the central peak signal (± 700bp from the center) was similar in WT and *Spo11KO* spermatocytes (Figure 2E, Table S4). Conversely, in the absence of MEI4 or IHO1, both HORMAD1 and SYCP3 signals at hotspot centers were reduced compared with the WT and *Spo11KO* signals (Figure 2C, 2E, Figures S4D-E, and Table S4). In *Mei4KO* and *Iho1KO* mice, we detected very few HORMAD1 or SYCP3 peaks overlapping with hotspots (4-5% of all peaks) (Tables S2 and S3). Moreover, HORMAD1 and SYCP3 were undetectable at hotspots in *Prdm9KO* spermatocytes (Figure 2C, Figure S4D). Altogether, these results suggest that the axis proteins HORMAD1 and SYCP3 bind at the hotspot center independently of DSB formation and thus presumably before break formation. Robust binding of HORMAD1 and SYCP3 to hotspots requires MEI4 and IHO1, but HORMAD1 and SYCP3 can also be partly recruited to hotspots through IHO1-and MEI4-independent modes.

### HORMAD1 and SYCP3 display distinct binding dynamics at CTCF and functional elements

HORMAD1 and SYCP3 ChIP-seq signal in WT spermatocytes also revealed peaks that localized outside hotspots (Figure 3A): 37% and 22% of HORMAD1 and SYCP3 peaks, respectively, overlapped with CTCF sites (Figure 3B, Tables S2 and S3). Moreover, a smaller subset of HORMAD1 and SYCP3 peaks overlapped with functional elements (FE) (Figure 3B), which we defined as the subset of gene regulatory elements that are enriched in H3K4me3 at zygonema and from which hotspots have been removed (see Star Methods). As some FE also contained CTCF sites, we defined three subcategories of sites: FE without CTCF (FE – CTCF), and CTCF sites with/without FE (CTCF + or –FE). Specifically, 13% and 8% of HORMAD1 and SYCP3 peaks were at FE (FE-CTCF and CTCF+FE), respectively (Figure 3B, Table S2 and S3). A small proportion of peaks (∼15% for both proteins) did not overlap with known genomic elements, and we named them ‘undefined’ (Un) (Figure 3B). We performed all signal intensity quantifications at CTCF-FE and at FE-CTCF. Heatmaps showed that HORMAD1 was enriched at the center of CTCF sites and correlated weakly with CTCF intensity (Figure 3C, Figure S5A). The SYCP3 signal was detectable on the browser and on the heatmap at strong CTCF sites, but was generally not above background noise (Figure 3A, 3C, S5C and S5E). At FE, HORMAD1 and SYCP3 enrichment was maximal at the FE peak center (Figure 3D) and was weakly correlated with H3K4me3 enrichment (Figure S5B). We then tested the functional dependencies for the enrichment at CTCF sites and FE.

**Figure 3.**
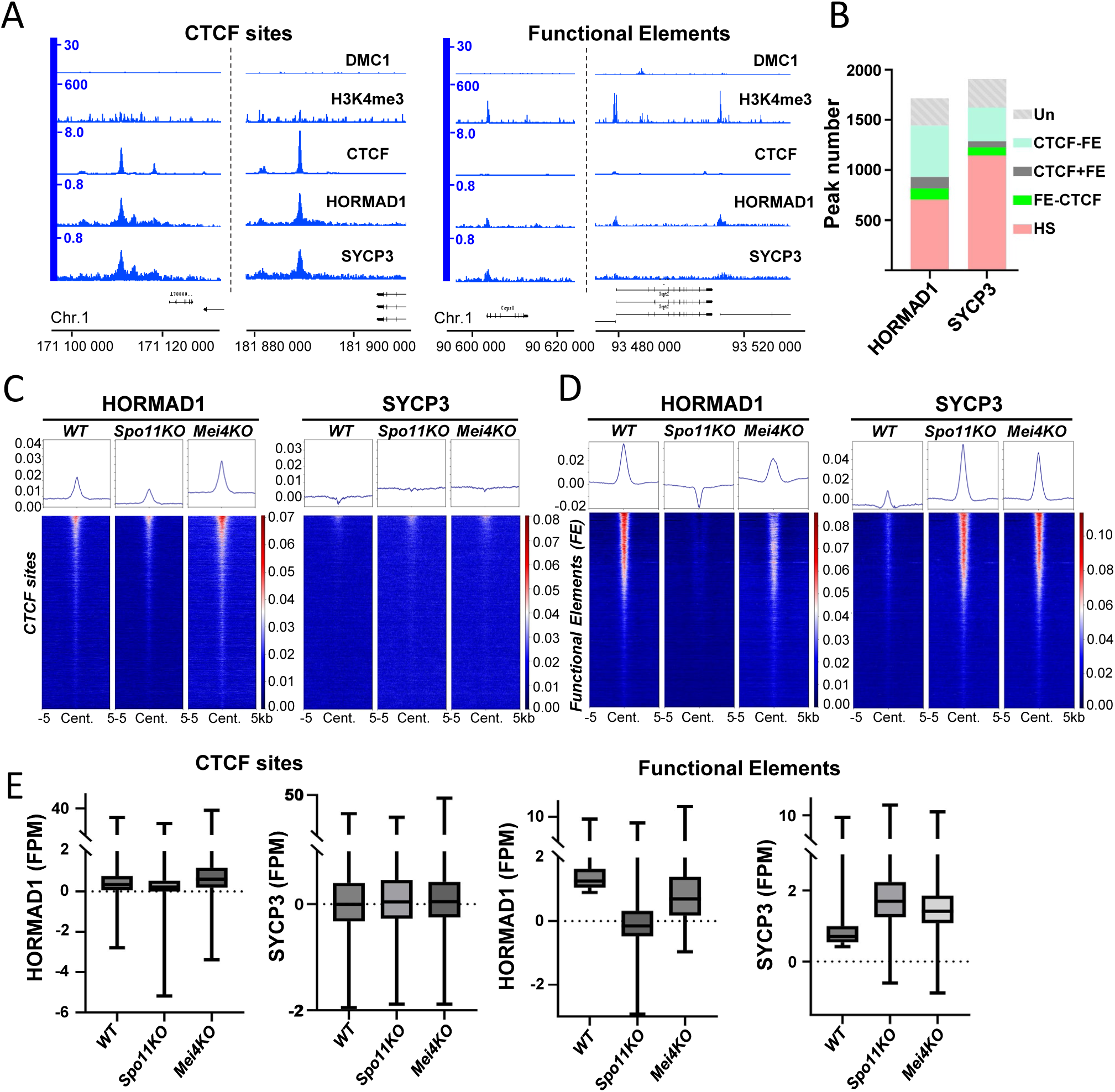
HORMAD1 and SYCP3 are enriched at CTCF and FE sites. **(A)** ChIP-seq read distribution of HORMAD1 and SYCP3, DMC1-SSDS ^31^, and H3K4me3 (zygonema) ^86^ at two representative CTCF sites and three representative FE in wild type (WT) spermatocytes. **(B)** Number of peaks at five genomic site types: hotspots (HS), FE-CTCF, CTCF+FE, CTCF-FE, and undefined sites (Un). **(C)** HORMAD1 and SYCP3 ChIP-seq signal in *WT*, *Spo11KO* and *Mei4KO* spermatocytes at CTCF (CTCF-FE) sites. CTCF signal was assessed by ChIP-seq in synchronized wild type (*B6)* testes. **(D)** HORMAD1 and SYCP3 ChIP-seq signal in *WT*, *Spo11KO* and *Mei4KO* spermatocytes at FE (FE-CTCF). H3K4me3 ChIP-seq signal intensity at FE was from ^86^. **(E)** HORMAD1 and SYCP3 ChIP-seq signal intensity at CTCF sites (left panels) and FE (right panels) in *wt*, *Spo11KO* and *Mei4KO* spermatocytes. The signal was measured at the 5000 strongest CTCF sites and FE.

At CTCF sites, we detected HORMAD1 and SYCP3 enrichment also in *Spo11KO* and *Mei4KO* spermatocytes with small variations, excepted for a marked increase of HORMAD1 intensity in *Mei4KO* (Figure 3C, 3E, Figures S5C, E, F). If HORMAD1 were limiting, this increase could be explained by the decreased HORMAD1 binding at hotspots in the *Mei4KO* strain. The weak SYCP3 signal at CTCF sites did not vary in the different genotypes tested (Table S4). At FE, HORMAD1 enrichment was lower in *Spo11KO* than in *Mei4KO* spermatocytes, also a potential indirect consequence of HORMAD1 occupancy at hotspots (Figure 3D, 3E, Figure S5C). Conversely, SYCP3 signal at FEs was increased in both *Spo11KO* and *Mei4KO* compared with WT spermatocytes, as shown by the higher percentage of SYCP3 peaks at FE (4% in WT *vs* 41% and 48% in *Spo11KO* and *Mei4KO*) and the median enrichment at FE (Figure 3D-E, Tables S3, S4, Figure S5C). This was accompanied by a reduced relative enrichment at CTCF sites in both *Spo11KO* and *Mei4KO* compared with WT, as shown by the ratios of the peaks overlapping with CTCF-FE and CTCF+FE (Figure S5D,tableS2, Material and Methods). This suggests changes in SYCP3 interaction with FE due to DSB formation, DSB repair, or downstream consequences of DSB repair, such as synapsis.

### IHO1, a DSB protein with axis protein features

IHO1 is essential for DSB activity ^4^. IHO1 forms a complex with REC114 and MEI4 ^19^ and localizes at early prophase as foci that mostly overlap with these two proteins ^4^. However, as axes extend and prophase progresses towards zygonema, IHO1 foci elongate and progressively extend along the unsynapsed axes, overlapping with HORMAD1 and SYCP3 ^4^. Conversely, MEI4 and REC114 remain as discrete foci along the unsynapsed axes ^17,18^. Importantly, IHO1 also interacts with the axis protein HORMAD1 ^4,50^.

We next analyzed IHO1 genomic localization. In WT spermatocytes, IHO1 localized at hotspots, CTCF sites, and weakly at FE (Figures 4A-B, Tables S2, S3). At hotspots, IHO1 showed a triple peak profile and the flanking peaks overlapped with HORMAD1 and SYCP3 peaks and with the end-resection profile (Figures 4A, 4C, Figure S6A). As observed for HORMAD1 and SYCP3, in the absence of SPO11, IHO1 formed a single central peak. However, IHO1 signal at hotspots was strongly reduced in *Mei4KO* mice and undetectable in *Prdm9KO* mice (Figure 4C). This was confirmed by the very low number of IHO1 peaks overlapping with hotspots in *Mei4KO* and *Prdm9KO* (Figure S6B, Tables S2, S3) and by the reduced median enrichment, signal intensity, and Cohen’s D values (Figure 4D, Figure S6C, Table S4). These data suggest that IHO1 recruitment at hotspots is PRDM9– and MEI4-dependent, but DSB formation-independent, and that upon DSB formation IHO1 interacts with resected DSB ends.

**Figure 4.**
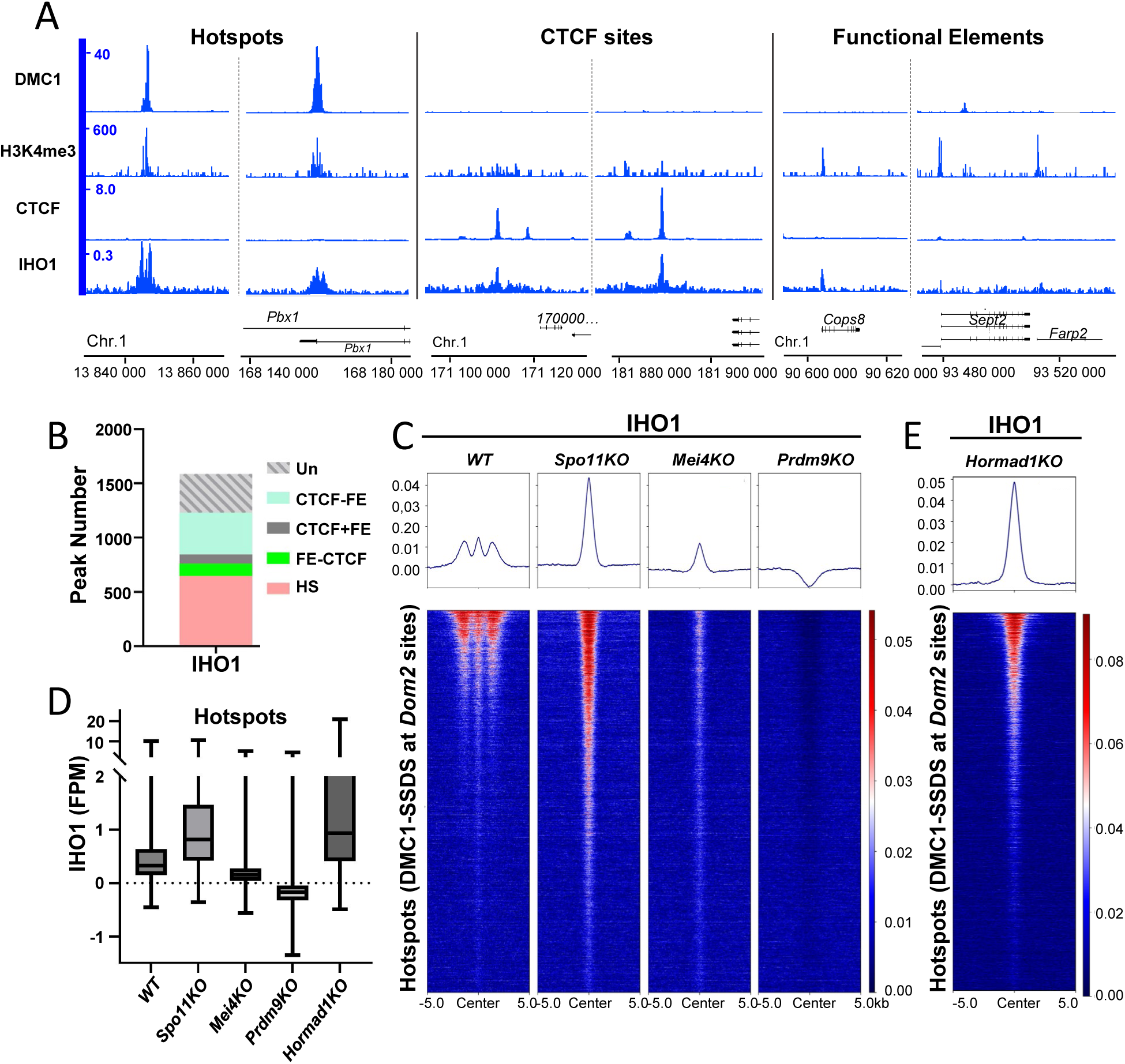
IHO1, a link between DSB proteins and axis proteins at hotspots. (**A**) ChIP-seq read distribution of IHO1, DMC1-SSDS ^31^, CTCF and H3K4me3 (zygonema) ^86^ at two representative hotspots and CTCF sites and at three representative FE in wild type (WT) spermatocytes. **(B)** Number of peaks at five genomic site types as in Figure 3B. **(C)** IHO1 ChIP-seq signal in *WT*, *Spo11KO, Mei4KO and Prdm9KO* spermatocytes at all *Dom2* hotspots ^31^. DMC1-SSDS signal intensity was from ^31^. **(D)** IHO1 ChIP-seq signal intensity at hotspots in *wt*, *Spo11KO*, *Mei4KO* and *Prdm9KO,* and *Hormad1KO* spermatocytes. The signal was measured at the 2000 strongest hotspots. **(E)** IHO1 ChIP-seq signal at all *Dom2* hotspots in *Hormad1KO* spermatocytes. DMC1 SSDS ChIP signal intensity was from ^31^.

In *Hormad1KO* mice, IHO1 signal was detectable but appeared as a single, strong peak, similar to the one observed in *Spo11KO* (Figure 4E). DSB activity is reduced by ∼4-fold in *Hormad1KO* ^47^; however, it is not known where the remaining breaks take place along the genome. Therefore, the loss of lateral peaks in these mutants can be interpreted in several ways: (i) the interaction of IHO1 at resected DSB ends (but not at the hotspot center) depends on HORMAD1; (ii) the 4-fold reduction of DSB formation at hotspots leads to a lateral peak signal reduction below the detection threshold; (iii) the DSBs generated in *Hormad1KO* mice are formed outside PRDM9-dependent hotspots. To test the third hypothesis, we determined DSB localization by DMC1-SSDS. In *Hormad1KO,* 95.2% of the DMC1-SSDS peaks localized within PRDM9-dependent hotspots, indicating no alteration in DSB localization control in the absence of HORMAD1 (Figure S6D). Despite the efficient hotspot binding by MEI4 and IHO1 detected by ChIP-seq in *Hormad1KO* mice (Figures 4E, S6E), at the cytological level, IHO1 (and MEI4) appear as weak foci that do not spread along the axis and/or are not axis-associated ^4^. Therefore, we re-evaluated the number of axis– and non-axis-associated MEI4 foci in *Hormad1KO* mice by setting a low detection threshold to detect weak foci. Although a subset of MEI4 foci was still localized on axes, in *Hormad1KO*, MEI4 foci were significantly more abundant outside the axes compared with WT (Figure S6F-G). These observations also showed that efficient MEI4 and IHO1 binding to hotspots was not sufficient for normal DSB activity and that HORMAD1 and/or axis-association was also required. Altogether, these binding features allowed us to propose a coherent pathway for the hierarchical recruitment of these proteins at hotspots where MEI4 promotes IHO1 binding, followed by axis association through the binding of HORMAD1 and SYCP3.

### Distinct dependencies between IHO1 and HORMAD1 at CTCF and FE sites

IHO1 and HORMAD1 showed a similar peak distribution in WT (41% for both are at hotspots and 24 and 30%, respectively, are at CTCF-FE) (Table S3). However, dependency between IHO1 and HORMAD1 was different at CTCF sites and FE. At CTCF sites, IHO1 enrichment depended on HORMAD1, while HORMAD1 was readily detectable in the absence of IHO1 (Figure 5A-B, Table S4). Conversely at FE, HORMAD1 intensity was reduced in *Iho1KO* (Figure 5C, Table S4). IHO1 intensity, which was weak at FE and with a smaller peak number (Tables S2 and S3), was not decreased in *Hormad1KO* (Figure S7A, Table S4). SYCP3 enrichment at CTCF sites (Figure S7B) and FE (Figure 5D, Table S4) did not depend on HORMAD1 or IHO1. SYCP3 increase at FE in *Iho1KO* (Figure 5D) was similar to the increase observed in the other DSB-defective mutants *Spo11KO* and *Mei4KO* (Figure 3D).

**Figure 5.**
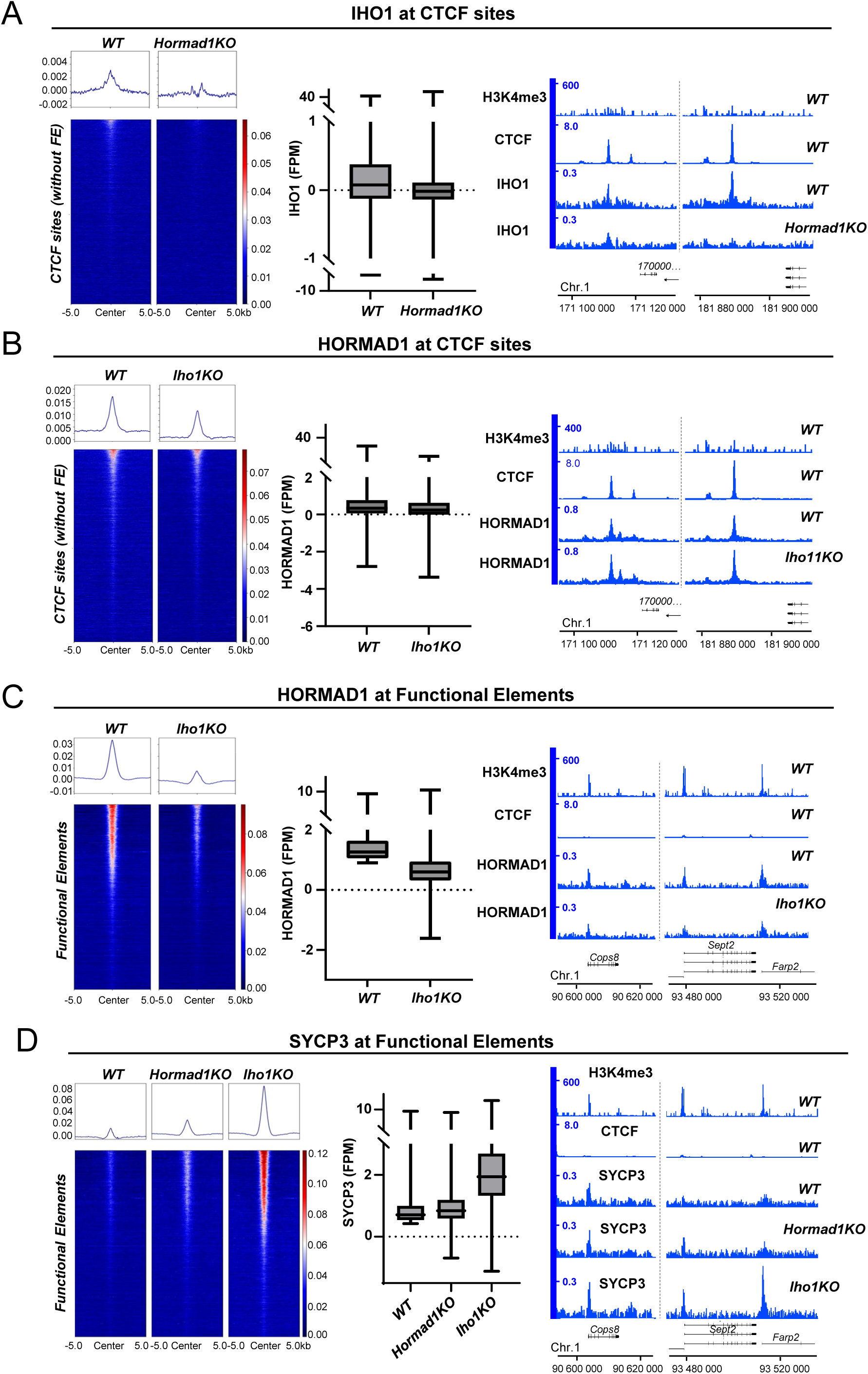
Opposing properties of HORMAD1 and SYCP3. (**A**) Left: IHO1 ChIP-seq signal at CTCF sites in wild type (WT) and in *Hormad1KO* spermatocytes. Middle: IHO1 ChIP-seq signal intensity at CTCF sites in WT and *Hormad1KO* spermatocytes. The signal was measured at the 5000 strongest CTCF sites. Right: ChIP-seq read distribution of IHO1, H3K4me3, and CTCF in WT and of IHO1 in *Hormad1KO* spermatocytes at two representative CTCF sites. **(B)** Left: HORMAD1 ChIP-seq signal at CTCF sites in WT and *Iho1KO* spermatocytes. Middle: HORMAD1 ChIP-seq signal intensity at CTCF sites in WT and *Iho1KO* spermatocytes. The signal was measured at the 5000 strongest CTCF sites. Right: ChIP-seq read distribution of HORMAD1, H3K4me3 (zygonema) ^86^ and CTCF (synchronized testes) in WT and of HORMAD1 in *Iho1KO* spermatocytes at two representative CTCF sites. **(C)** Left: HORMAD1 ChIP-seq signal at FE in WT and *Iho1KO* spermatocytes. Middle: HORMAD1 ChIP-seq signal intensity at FE in WT and *Iho1KO* spermatocytes. The signal was measured at the 5000 strongest FE. Right: ChIP-seq read distribution of HORMAD1, H3K4me3 ^86^ and CTCF in WT and of HORMAD1 in *Iho1KO* spermatocytes at three representative FE. **(D)** Left: SYCP3 ChIP-seq signal at FE in *WT*, *Hormad1KO* and *Iho1KO* spermatocytes. Middle: SYCP3 ChIP-seq signal intensity at FE in *WT, Hormad1KO* and *Iho1KO* spermatocytes. The signal was measured at the 5000 strongest FE. Right: ChIP-seq read distribution of SYCP3, H3K4me3 ^86^ and CTCF in WT*, Hormad1KO* and of SYCP3 in *Iho1KO* spermatocytes at three representative FE.

These functional analyses highlighted unexpected dynamic interactions and binding principles of axis components. First, unlike SYCP3, IHO1 and HORMAD1 were relatively more bound to CTCF sites than FE in *Spo11KO* and *Mei4KO* (Table S2). Second, the dependency between IHO1 and HORMAD1 was reversed at CTCF and FE sites: IHO1 was required for HORMAD1 binding at FE but not at CTCF sites (Figure 5B-C). Third, SYCP3 binding to CTCF sites and FE was HORMAD1-independent. Fourth, we detect a systematic increase of SYCP3, but not HORMAD1, enrichment at FE in the absence of DSB activity (Figure 3D, 5D).

### Implications for DSB activity in the absence of PRDM9

In the absence of PRDM9, most DSB activity takes place at promoters and enhancers, called default sites ^36^. It was shown that 44% of these sites overlap with annotated genes ^36^ and 29% overlap with FE, as defined in our study. Therefore, we wondered whether HORMAD1 and SYCP3 enrichment at FE in WT highlights some features relevant to DSB formation in *Prdm9KO* mice. In fact, we observed a weak positive correlation between HORMAD1 enrichment at default sites in WT mice and DSB level (DMC1-SSDS) in *Prdm9KO* mice (Figure 6A). We observed a similar correlation between SYCP3 enrichment at default sites (in *Spo11KO*) and DMC1-SSDS in *Prdm9KO* mice (Figure 6A). This suggests that when PRDM9 is absent, axis protein enrichment at H3K4me3 enriched sites contributes to DSB activity, but that other factors also contribute to default hotspot strengths.

**Figure 6.**
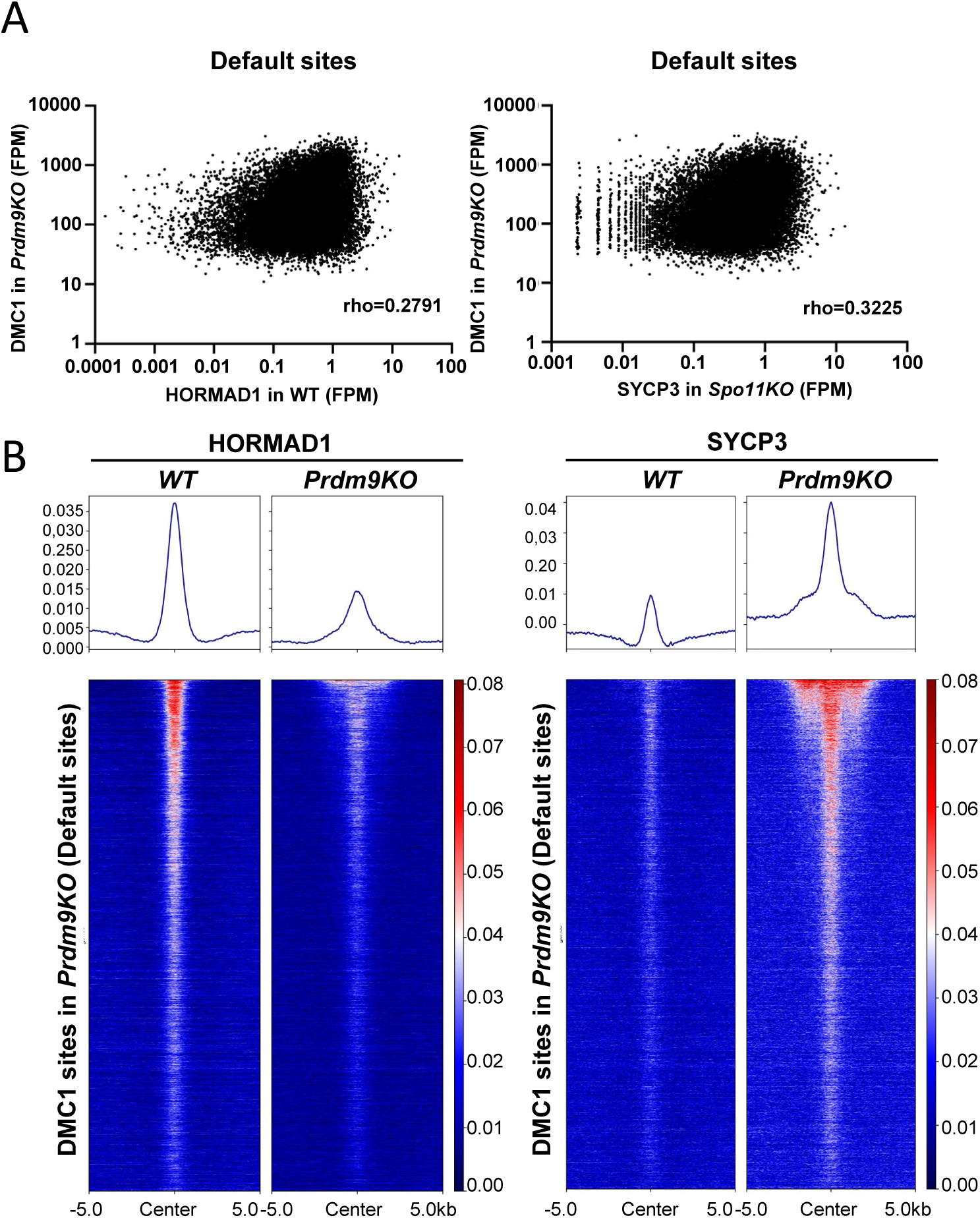
Default sites are enriched for axis proteins. **(A)** HORMAD1 (left) and SYCP3 (right) signal intensity (FPM) in wild type (WT) and *Spo11KO* compared with the DMC1 signal intensity in *Prdm9KO* strains ^36^ at default sites. roh: Spearman correlation coefficient. **(B)** HORMAD1 (left) and SYCP3 (right) ChIP-seq signal at default sites in WT and *Prdm9KO*. DMC1 signal intensity was from ^36^.

Next, we analyzed MEI4, HORMAD1 and SYCP3 recruitment at default sites in *Prdm9KO*. We did not detect MEI4 at default sites in *Prdm9KO* mice (Figure S1K). Conversely, HORMAD1 and SYCP3 signal distribution showed a larger peak compared to WT (Figure 6B). On the heat map, SYCP3 signal showed two peaks flanking the center of the default sites at the strongest default sites, based on DMC1-SSDS intensity. We propose that, similarly to what we observed in WT spermatocytes at PRDM9-dependent hotspots (Figure 2), this signal spreading away from the default site center may be correlated to DSB end-resection. Thus, similarly to PRDM9-dependent hotspots, SYCP3 (and potentially HORMAD1) seems to be recruited near the end of resection tracts at DSBs formed in the absence of PRDM9.^36^

### DSB hotspots contact CTCF sites

We previously showed that PRDM9 ChIP yielded a faint, but distinct signal at CTCF sites and some FE ^31^. In addition, a benzonase sensitive co-immunoprecipitation experiment to assess PRDM9-CTCF interaction led us to conclude that hotspots might be in physical proximity of CTCF sites and that this interaction may occur over long distances, likely involving chromatin ^31^. Here, we performed a CTCF ChIP-seq experiment using leptotene/zygotene spermatocytes from synchronized WT testes. We detected 28134 peaks and found that CTCF also yielded a faint, but specific signal at hotspots (Figure 7A). Unlike PRDM9, which shows a single sharp peak at CTCF sites ^31^, the CTCF enrichment at hotspots yielded a broad peak with a bimodal distribution (Figure 7B-C) comparable to that of IHO1 and axis proteins (see Figure 2), showing the presence of CTCF on resected DSB ends. Indeed, like for IHO1, HORMAD1 and SYCP3, the lateral peaks were lost in *Spo11KO* (Figure 7B) and the CTCF signal intensity at hotspots correlated with that of HORMAD1 and SYCP3 (Figure 7D).

**Figure 7.**
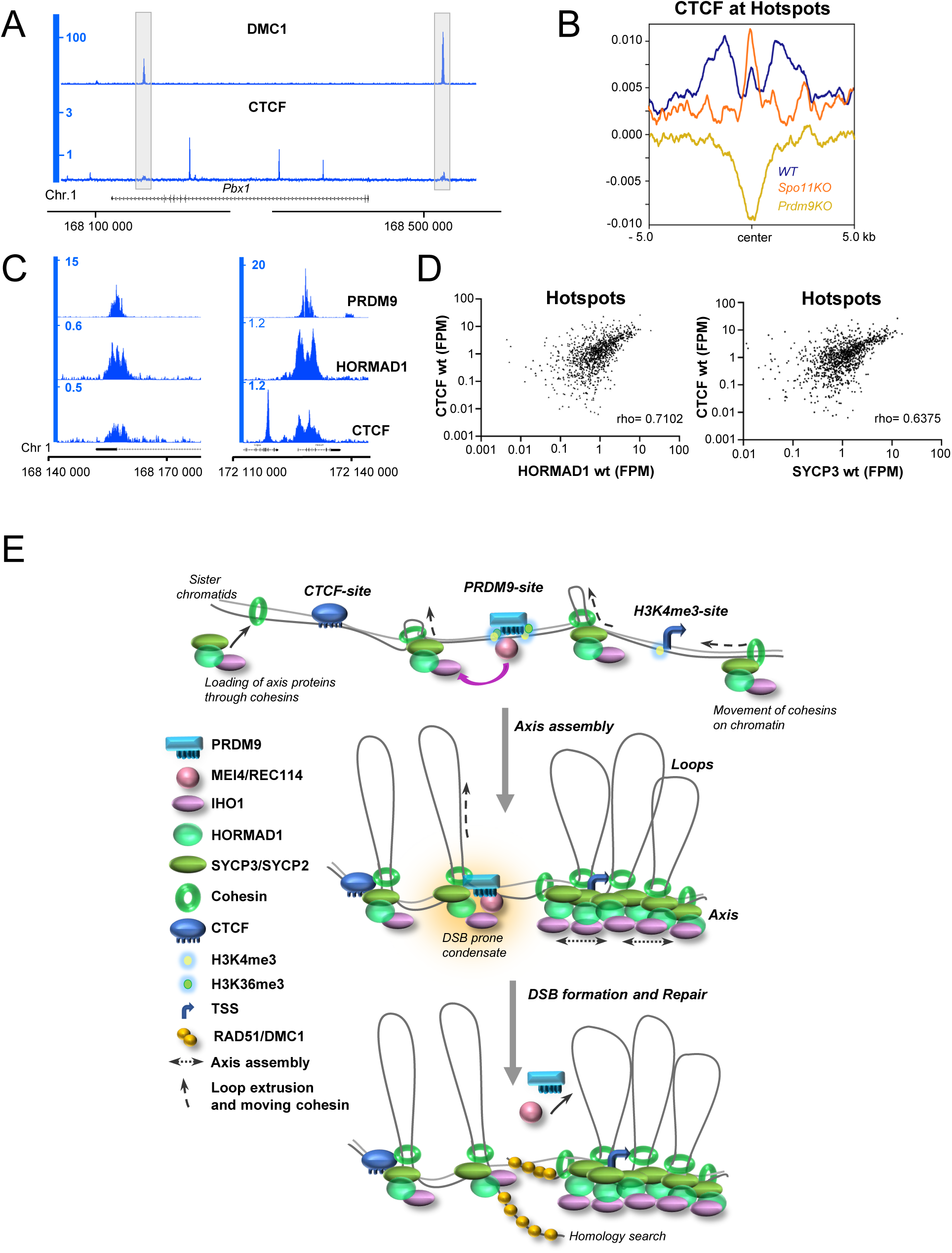
DSB hotspots contact CTCF sites. (**A**) ChIP-seq read distribution of CTCF and of DMC1-SSDS at two representative hotspots (Chr1; 168,100,000-168,500,000) in wild type (WT) mice. Gray boxes highlight non-canonical CTCF signals coinciding with DSB sites. **(B)** Averaged enrichment profile of CTCF ChIP-seq at hotspots in WT (blue), *Spo11KO* (orange) and *Prdm9KO* (yellow) spermatocytes at *Dom2* hotspots. **(C)** ChIP-seq read distribution of PRDM9, HORMAD1, and CTCF in WT spermatocytes at two representative hotspots. **(D)** HORMAD1 (left) and SYCP3 (right) signal intensity (FPM) compared with the CTCF signal intensity at *Dom2* hotspots in WT spermatocytes. roh: Spearman correlation coefficient. **(E)** At meiosis entry, cohesins promote axis proteins loading on chromatin. Cohesin complexes can potentially exist in two types: extruding complexes and cohesive complexes. Cohesin/axis complexes move along chromatin and are stabilized at CTCF sites, PRDM9 sites and other functional elements enriched in H3K4me3 that act as barriers. Loop basis may be stabilized by the filamentous assembly and DNA bridging properties of axis proteins. The localization of PRDM9-sites at loop bases favors the assembly of DSB-formation protein condensates. DSBs are formed, resected and repaired by homologous recombination in the context of axis.

## DISCUSSION

In this work, we present information on the interplay between essential components of the meiotic DSB machinery and chromosome axis formation in mammals. We propose that three genomic site types contribute to the loop/axis organization of meiotic chromosomes: *Prdm9*-dependent hotspots, CTCF binding sites, and H3K4me3-enriched sites. Based on the enrichment of MEI4, IHO1, HORMAD1 and SYCP3 at these genomic sites and their functional dependencies, we identified key properties of the chromosome organization at early prophase, and we propose a model on how this organization may be put in place.

### Interaction between DSB sites and axis

A widely conserved feature of meiotic DSB repair is that it takes places in the context of the meiotic chromosome axis. The axis appears to be a platform and a structure to ensure several regulations of DSB repair, such as the homolog bias and the crossover/non-crossover choice ^6^. To allow DSB repair regulation by axis components, one molecular strategy adopted in several species is to catalyze DSBs on axis-associated DNA sequences ^50^. This interplay between axis and DSB sites has been extensively studied in *S. cerevisiae*. It has been shown that at prophase onset, chromosomes are organized as arrays of loops anchored to the chromosome axes. At this stage, the essential DSB accessory factors (Rec114, Mei4, Mer2) colocalize on the axis, and the potential DSB sites, which are mainly promoters, are in the loops ^9,10^. DSB sites are thought to be tethered to the axis by interaction between Spp1 (bound to DSB sites) and Mer2 ^8,11^. In this scenario, Mer2 is required for Mei4 interaction with DSB sites.

In mice (and some other mammals), PRDM9 is implicated in DSB control. We propose that PRDM9 uses a different molecular strategy to reach the same objective: to localize DSB sites on the axis. We found that in mouse spermatocytes, MEI4 is enriched at PRDM9-dependent DSB sites in an IHO1 (the mouse Mer2 ortholog)-independent manner, and IHO1 enrichment at these sites is MEI4-dependent (Figures 1C, 1E, 4C). Therefore, unlike in *S. cerevisiae*, the interaction between DSB sites and axis is mainly specified by MEI4, and not IHO1. We propose that PRDM9-bound sites are anchored to axes by the interaction between MEI4, IHO1 and axis-components (Figure 7E). Indeed, at PRDM9-bound sites, we also showed that HORMAD1 and SYCP3 are recruited mainly in a MEI4– and IHO1-dependent manner (Figures 2C, 2E). We also detected a low level of MEI4-independent enrichment of IHO1, HORMAD1 and SYCP3 at DSB sites, suggesting an alternative pathway (Figure 2C, 2E, 4C). Our study did not allow determining the kinetics and spatial organization of these interactions. We could hypothesize that HORMAD1 and SYCP3 are pre-assembled on chromatin at genomic sites where we detected their enrichment (i.e. CTCF and H3K4me3 sites) and where they may be recruited by interaction with cohesins (see below “Building the axis). Then, this axis-associated pre-assembled complex would interact indirectly with MEI4 and/or REC114 at DSB sites. This could be mediated by IHO1 that has the potential to bind to HORMAD1 ^4,50^ and form a complex with MEI4 and REC114 (RMI complex)^19^ (Figure 7E).

Thus, besides the different roles of IHO1/Mer2 and MEI4/Mei4, the process of linking DSBs to axes is conceptually similar in mouse and *S. cerevisiae*. A similar principle seems to govern interactions between DSB and axis sites in *Schizosaccharomyces pombe*, where the axis components Rec10 (SYCP2/Red1 ortholog) and Hop1 (HORMAD1 ortholog) are enriched at DSB sites ^51–53^. Rec15 (IHO1/Mer2 ortholog) also plays a central role in these interactions, providing a link between axis and DSB sites, and its interaction with DSB sites is mutually dependent with Rec24 (the MEI4/Mei4 ortholog) ^52,53^. In mice, how MEI4 is recruited to PRDM9-bound sites remains to be understood. As PRDM9 methyltransferase activity is required (Figure S3B), histone modifications and/or a reader of such modifications might be involved. The reader ZCWPW1 is not required for DSB formation ^54–56^, but it could be interesting to test whether the paralog ZCWPW2 is involved. The putative RMI complex may be stabilized upon interaction with axis components and its ability to form DNA-driven condensates also might be enhanced ^20,21,50^. In this context, we predict that the cytologically detectable MEI4/REC114/IHO1 foci (∼300 foci) are associated with DSB sites where these proteins can activate the SPO11/TOPOVIL catalytic complex partly through the REC114 –TOPOVIBL interaction ^23^. Two other proteins, ANKRD31^57,58^ and MEI1 ^32,59^, also might contribute to this DSB activation in an unknown manner. Disrupting the interaction of DSB proteins with the axis, such as in the *Hormad1* mutant or the *Iho1* mutant that cannot interact with HORMAD1, leads to a decrease in DSB activity despite the assembly of the DSB proteins on chromatin ^4,17,47,50^(Figure S6F-G). Therefore, axis-binding of RMI complexes (as opposed to chromatin-binding off-axis) is necessary for the efficient activation of the catalytic DSB complex. The connection between DSB activity and axis binding gains particular significance during homolog synapsis, where HORMAD1, IHO1 and its partners MEI4 and REC114 are evicted from the axes, contributing to turning off DSB activity ^60,61^.

Interestingly, in the absence of PRDM9, DSBs form at default sites, including promoters and enhancers ^36^. In this context, DSB site-axis interactions might be promoted through a mechanism similar to what described in *S. cerevisiae*: IHO1, localized at axis-sites with HORMAD1 and other axis components, might be driving the interaction of DSB sites to the axis. This would require a specific interaction of IHO1 with these default sites, for instance with CXXC1 (the ortholog of *S. cerevisiae* Spp1), which interacts with IHO1 in a yeast two-hybrid assay ^62^. In this scenario, MEI4 associated with IHO1 will indirectly be located in the proximity of DSB sites. We did not detect significant enrichment of MEI4 and IHO1 at default sites in *Prdm9KO* spermatocytes. This could be explained by limitations of the ChIP approach due low signal for distant and indirect protein-DNA interactions and low average occupancy in cell populations.

### Building the axis

The interaction between DSB sites and axis, which we estimated based on the number of MEI4 foci, is predicted to represent only a minority (<10%) of axis-association sites along chromosomes, as several hundred loops are expected to organize each mouse chromosome.

The formation of axes and the associated chromatin loop arrays might be largely driven by cohesin rings that move along chromatin and extrude chromatin loops and/or promote loop formation without ATP-driven motor activity. This should result in the stacking of cohesin rings in a linear configuration, stabilized by axis proteins (SYCP2 and SYCP3) (Figure 7E). The two cohesin complexes, involving the kleisin subunit REC8 or RAD21L, interact and colocalize with the structural axis proteins SYCP2, SYCP3 and HORMAD1 ^15^ (Figure 7E). REC8, but not RAD21L, is required for sister chromatid cohesion ^63^, and it is unknown whether both cohesin complexes mediate loop extrusion. As *S. cerevisiae* Red1 interacts with Rec8 ^64^, it is tempting to speculate that SYCP2 (the mouse Red1 ortholog) also interacts with cohesins. SYCP2 oligomerizes with SYCP3, forming bundles *in vitro* ^40^ that may link adjacent cohesins in the axis. Moreover, axis compaction may be provided by the DNA bridging property of SYCP3 ^65^. Furthermore, SYCP2 may recruit HORMAD1 or stabilize its interaction with cohesins ^15^. The cohesin complexes can be stalled on chromatin at specific genomic locations, such as CTCF sites and promoters. Indeed, it has been shown that REC8 and RAD21L are enriched at CTCF sites and promoters in mouse spermatocytes ^27^ (Figure 7E). Our identification of SYCP3 and HORMAD1 enrichment at CTCF and H3K4me3-enriched sites (Figure 3) is consistent with the stalling of cohesins and associated axis proteins at these sites. that PRDM9 binding sites could also act as cohesin anchor sites and/or barriers to loop extrusion. As described above (see “Interaction between DSB sites and axis”), anchoring of PRDM9-bound sites to axes is predicted to take place concomitantly with axis formation. In agreement, in our previous and current studies, we detected contacts predicted by this organization: by immunoprecipitation and crosslinking ChIP, CTCF and PRDM9 show slight enrichment at each other sites ^31^(Figures 7A-D). PRDM9 also co-immunoprecipitates with RAD21L and REC8^35^. Unlike studies in yeast ^66–68^, the recent mouse Hi-C analyses detected only few specific genomic contact points in prophase spermatocytes at leptonema and zygonema ^24,27,29,69^. This could be due to the complexity of contact regions that might limit their detectability in a cell population. When cohesins are depleted, such as in *Stag3KO* or *Rec8 Rad21L double KO*, SYCP3 and HORMAD1 assembly is impaired, MEI4 foci are poorly detectable, and DSB activity is strongly reduced ^17,35,63,70,71^. In *Sycp3KO* mice, cohesins assemble on the axis^46^, HORMAD1 shows a punctate staining on the axis where MEI4 foci localize, and DSBs form (Figure S7). Mice with reduced CTCF levels show no defect in meiotic prophase (73), suggesting that DSB formation and repair are globally not affected ^72^, suggesting that DSB formation and repair are globally not affected. It is possible that the change in loop organization is relatively modest because CTCF insulator activity is low in early meiotic prophase ^24,27,29,69^. Other boundary elements may contribute to stabilize the cohesin complexes. In addition, as loop sizes have been shown to increase during meiotic prophase ^24,29,69^, fixed boundaries might not be required and the loop-axis organization might be highly dynamic.

### Axis proteins are recruited at processed DSB ends

After DSB formation, in mouse spermatocytes, DSB ends are resected by the coordinated action of MRN, CTIP, ExoI and BLM/DNA2 that leads to the resection of 0.3 to 2.0Kb on both ends ^48,49,73^. Here, we discovered that SYCP3, HORMAD1 and IHO1 localize to resection tract ends (Figure 7E). How they are recruited to DSB ends remains to be determined. As HORMAD1 and SYCP3 interact with cohesins in meiotic prophase and cohesins, in somatic cells, have been detected at DSB ends ^74–76^, one possible scenario is the recruitment of HORMAD1 and SYCP3 by cohesins at meiotic resected DSB ends. Alternatively, or in addition, HORMAD1 and/or IHO1 may interact with the MRN (MRE11/NBS1/RAD50) complex involved in end resection, as proposed in *S. cerevisiae* ^77,78^, *Caenorhabditis elegans* ^79^ and *Arabidopsis thaliana*^80^. It has been suggested that the two ends of each DSB have distinct properties: one engaged in interaction with the sister-chromatid and potentially the axis, and the other engaged in homology search ^81^. As the signal recovered by ChIP is a population average, we do not know whether both ends are loaded with the axis proteins at a single DSB. As HORMAD1 is displaced from synapsed axes during zygonema ^41,42^, it will also be interesting to examine the interaction dynamics of these axis proteins during DSB repair from zygonema to pachynema.

The loop-axis organization of meiotic chromosomes is expected to have additional regulatory functions for DSB activity. Indeed, the axis length (and the predicted number of loops) correlates with the number DSB repair foci in female and male human meiosis ^82^. Also, in the pseudo-autosomal region of mouse sex chromosomes, shorter loops correlate with higher DSB activity ^83^. Therefore, loops may be functional units to control the DSB potential and DSB activity might be regulated by controlling the loop size, thus independently of the genome size ^9,84^. It has been also proposed that loops provide cis-regulation of DSB activity via Tel1, the *S. cerevisiae* ATM ortholog, along chromosomes ^85^.

### Limitations of the study

The data we obtained reveals multiple and complex protein-chromatin interactions at a large number of sites in the genome. Thus, several combinations of these interactions in single-cells could account for our population average view using ChIP. We do not think that single cell ChIP approach can be performed at this stage. However, high-throughput and high-resolution imaging approaches could certainly add complementary information. In addition, as we know from cytological analyses, and as we detect from the analysis of several mutants, the molecular steps that we have analysed are highly dynamic. We are therefore capturing an average of chromatin organization before, at and just after DSB formation. Although we have used highly purified and staged spermatocytes samples, additional approaches could be envisioned for a more precise timing of events.

## ACKNOWLEGDEMENTS

We thank the BioCampus Montpellier facilities for providing excellent technical support: the Reseau des Animaleries de Montpellier (RAM) for animal care, and Montpellier Resources Imagerie (MRI) for microscopy. We thank Sylvain Barrier and Aubin Thomas for providing help with the cluster and we are grateful to the Genotoul bioinformatics platform, Toulouse Midi-Pyrenees, for providing storage resources. We thank Callum Burnard for suggesting the use of Cohen’s D, and Alice Libri for help on *Sycp3KO* cytological analyses. We thank Giacomo Cavalli, Frédéric Baudat, Mathilde Grelon and Denise Zickler for critical reading of the manuscript. We also thank Marion Helsmoortel for technical help and Frédéric Baudat, Thomas Robert, Romain Koszul, Helene Bordelet and Tom Sexton for fruitful discussions.

## AUTHOR CONTRIBUTIONS

Conceptualization: CG, BdM, Methodology: CG, MB, BdM, AT, Experiments: MB (ChIP, Cytology, Imaging), CG (ChIP), LG (cytology), CB (cytology); Analysis: MB (bioinformatics), BdM (quantifications, correlations), CG (cytology), Funding acquisition: BdM, Supervision: CG, BdM, Writing – original draft: CG, BdM, Writing – review & editing: CG, BdM, MB, AT.

## MATERIALS and METHODS

### Resource Availability

#### Lead contact

Further information and requests for resources and reagents should be directed to the lead contacts, Corinne Grey

#### Materials availability

All unique reagents generated in this study are available from the lead contacts with a completed Materials Transfer Agreement.

#### Data and code availability

- All ChIP-seq and processed data generated in this study are publicly available at GEO with the accession number GSE262343, as the date of publication. All Imaging data generated in this study are acessible at Mendeley Data database with DOI: 10.17632/wd9367jyvs.4
- This paper does not report original code.
- Any additional information required to reanalyse the data reported in this paper is available from the lead contacts upon request.

#### Experimental Model and Study participant Details

The following mouse strains were used: C57BL/6JOlaHsd (*B6*)(referred to as wild type, WT), B10.MOLSGR(A)-(D17Mit58-D17Jcs11)/Bdm (*RJ2*) ^87^, B6;129P2-Prdm9tm1Ymat/J (*Prdm9KO*)^88^, Spo11tm1Mjn (*Spo11KO*)^33^, C57BL/6J-Tg(RP23-159N6*)23Bdm (*B6-Tg(YF)*)^37^. Mei4tm1Bdm (*Mei4KO*)^3^, Sycp3tm1Hoog (*Sycp3KO*)^89^, Hormad1tm1.2Atot (*Hormad1KO*)^47^, and Iho1tm1.2Atot (*Iho1KO*)^4^. All experiments were carried out on juvenile male animals according to the CNRS guidelines and were approved by the ethics committee on live animals (project CE-LR-0812 and 1295, ethical approval APAFIS#20218-2019091211174938v2). All animals for ChIP where used at 8dpi (see synchronization protocol below). For MEI4 or DMC1/RPA2 foci count, animals were used at 8dpi (*Spo11KO, Prdm9KO, Hormad1KO*), at 12dpp (*Iho1KO*) or at 14dpp (*Sycp3KO)*. The phenotypes of the various mutant strains (DSB formation, axis formation and synapsis, stage of arrest) are specified in Table S5.

## Method details

### Synchronization of spermatogenesis

Synchronization was performed as described in ^90^ and adapted in ^91^. Briefly, 1.5dpp male pups were treated with a retinoic acid inhibitor (WIN 18,446, Tocris, Biotechne # 4736) (100µg/g of body weight every 22-24h) to allow the accumulation of B type spermatogonia. After 7-10 days of treatment, a single dose of retinoic acid (50µg in 10µL of dimethyl sulfoxide) (Sigma-Aldrich, R2625-50MG) was intraperitoneally injected to trigger entry in meiosis. Testes were harvested exactly eight days after this injection. At this time point, 85-100% of SYCP3-positive cells corresponded to spermatocytes in the leptotene or early/mid zygotene stage (Table S1). Staging was assessed by SYCP3, SYCP1 and γH2AX staining (except in spermatocytes from *Sycp3KO* mice where HORMAD1 was used instead of SYCP3) on spermatocyte spreads using a small portion of testis tissue. The remaining testis tissue was processed for ChIP.

### Antibodies

Guinea pig anti-SYCP3 ^87^, rabbit anti-SYCP1 (Abcam, 15090), rabbit anti-DMC1 (Santa Cruz, H100), anti-MEI4 ^3^, and mouse monoclonal anti-phosphorylated histone H2AX (Ser139) (γH2AX) (Millipore, 05–636) antibodies were used for immunostaining. For DMC1 ChIP-SSDS, a goat anti-DMC1 antibody (Santa Cruz, C-20) was used. For conventional ChIP experiments, rabbit anti-PRDM9 ^31^, rabbit anti-MEI4 ^3^, rabbit anti-IHO1 ^4^, rabbit anti-HORMAD1 ^42^, rabbit anti-SYCP3 (Abcam, ab15093), and rabbit anti-CTCF (Abcam, ab128873) antibodies were used.

### Spermatocyte spreading and Immunostaining

Spreads of spermatocyte nuclei were prepared with the dry down technique, as described ^92^, and immunostaining was performed as described ^87^. Staging criteria were as follows: pre-leptotene nuclei had weak SYCP3 nuclear signal and no or very weak γH2AX signal; leptotene nuclei were γH2AX-positive and SYCP1-negative; early/mid zygotene nuclei had less than five or nine fully synapsed chromosomes respectively; late zygotene had nine or more fully synapsed chromosomes; and pachytene cells had all chromosomes fully synapsed, but for the sex chromosomes. The following antibodies were used: guinea-pig anti-SYCP3 (1:500), anti-SYCP1 (1:400) and anti-γH2AX (1:10,000).

### Microscopy

Widefield images were acquired using a Zeiss Axioimager 100X Plan Apochromat 1.4 NA oil objective and a Zeiss CCD Axiocam Mrm 1.4 MP monochrome camera (1388 x 1040 pixels, 6.45µm pixel size).

### Image analysis

For quantification, images underwent deconvolution using Huygens Professional version 22.10 (Scientific Volume Imaging). All image analyses were performed using Fiji/ImageJ 1.53t 98. The “MeiQuant” macro was used for focus counting and intensity measurements ^93^. Briefly, single nuclei were cropped manually. Foci were detected using the Find Maxima function. On-axis and off-axis foci were distinguished on the basis of their localization within and outside a binary mask, respectively. This region of interest was drawn using an automatic SYCP3 axis protein staining threshold. The same mask was used for measuring focus intensity. Each focus was first defined as on– or off-axis, and then the pixel with maximum intensity was automatically detected using the Find Maxima function. Statistical analyses were performed with GraphPad Prism 9 and the nonparametric Mann-Whitney test to compare the number of foci and focus intensity.

### Chromatin immunoprecipitation (ChIP) and library preparation

ChIP experiments were performed with the ChIP-IT High Sensitivity Kit (Active Motif, 53040). Briefly, de-capsulated testes from two or three synchronized mice (see above) were homogenized and fixed at the same time in fixation solution for 15min. After quenching, tissues were homogenized, and cell suspensions prepared by filtering through a 40µm cell strainer. Cells were washed twice with ice-cold 1x PBS, and chromatin was extracted, sonicated and immunoprecipitated according to the manufacturers’ instructions. 30-40µg of chromatin was used per immunoprecipitation. The following antibodies (amount) were used: affinity purified rabbit anti-PRDM9 (4µg), affinity purified rabbit anti-MEI4 (4µg), affinity purified rabbit anti-HORMAD1 (4µg), affinity purified rabbit anti-IHO1 (4µg), rabbit anti-SYCP3 (4µg), and rabbit anti-CTCF (4 µg). Libraries were prepared using the Next Gen DNA Library kit from Active Motif (53216) following the manufacturer’s instructions. Sequencing was performed on a Novaseq sequencing machine using the paired end 150bp mode.

### DMC1-SSDS (DMC1-Single Strand DNA Sequencing)

After crosslinking whole testes, DMC1-ChIP was performed as in ^31^. ssDNA was enriched during library preparation as in ^31^. Testes from two synchronized mice (WT and *Hormad1KO* littermates) were used for each replicate. Sequencing was performed on a Novaseq sequencing machine using the paired end 150bp mode.

## Quantifications and statistical analysis

### Detection of DMC1-SSDS peaks

Raw reads were processed using the hotSSDS and hotSSDS-extra Nextflow pipelines ^94^. Briefly, the main pipeline steps included raw read quality control and trimming (removal of adapter sequences, low-quality reads and extra bases) and mapping to the UCSC mouse genome assembly build GRCm38/mm10. Single stranded derived fragments were then identified from mapped reads using a previously published method ^36,95^ and peaks were detected in Type-1 fragments (high confidence ssDNA). To control reproducibility and assess replicate consistency, the Irreproducible Discovery Rate (IDR) method ^96^ was used, following the ENCODE procedure (https://github.com/ENCODE-DCC/chip-seq-pipeline2). The “regionPeak” peak type parameter and default p-value thresholds were used. Briefly, this method performs relaxed peak calling for each of the two replicates (truerep), the pooled dataset (poolrep), and pseudo-replicates that are artificially generated by randomly sampling half of the reads twice, for each replicate and the pooled datasets. Both control and *Hormad1KO* datasets passed the IDR statistics criteria for the two scores (below 2). By default, the pipeline gives the poolrep as primary output, but for this study the truerep peak datasets were considered. Lastly, peak centring and strength calculation were computed using a previously published method ^95^.

### Read alignment and detection of ChIP-seq peaks

After quality control, ChIP-seq reads were trimmed to 100bp and filtered to keep the sequencing read quality Phred score > 28. Reads were then mapped to the UCSC mouse genome assembly build GRCm38/mm10 using Bowtie 2 (version 2.3.2) with the following parameters: –N 1 –I 100 –X 1000 –no-mixed –no-discordant. Then, only not duplicated and uniquely mapped reads were kept for the subsequent analysis. To identify the highly reproducible enriched regions from the filtered aligned fragments between replicate samples, the IDR methodology was used, as described in the ENCODE and modENCODE projects ^97^. This method allows testing the reproducibility within and between replicates by using the IDR statistics. Briefly, peak consistency is evaluated between (i) true replicates, (ii) pooled pseudo-replicates and (iii) self-pseudo-replicates. Following their pipeline, peak calling was performed on true replicates, pooled pseudo-replicates and self-pseudo-replicates with a relaxed p-value. Therefore, MACS2 (version 2.2.6) was run with the following parameters: –-format BAMPE –p value 0.01 as advised by the authors, and a negative control was included (the corresponding knockout, except for CTCF, where input was used) to reduce background noise. Then, IDR analyses were performed, and after comparing all obtained peak datasets, replicates with an IDR rescue and self-consistency ratio below 4 were considered to be reproducible. The final peak datasets were generated by taking the top N peaks from true replicates below the IDR threshold of 0.05, as recommended by the authors. If IDR conditions were not satisfied, replicates were considered not reproducible, and thus only one replicate was considered: the one with the highest percentage of peaks that passed the IDR threshold. The final peak dataset was generated by taking the top N peaks from pseudo-replicates below the IDR threshold of 0.01, as recommended by the authors ^97^. CTCF ChIP-seq yielded 32676 peaks. We filtered the overlap with FE and obtained a pool of 28134 CTCF-FE and 4542 CTCF+FE peaks. For all signal intensity quantifications at CTCF sites we used the subpopulation CTCF-FE unless otherwise noted.

### Peak classification for ChIP-seq

Final peak datasets were annotated with annotatePeaks.pl from the HOMER suite (version 4.9.1). Overlap were determined with the bedtools suite (version 2.26.0) using the module intersect and 1bp overlap. The relative enrichment of IHO1, HORMAD1 and SYCP3 at CTCF vs FE can be evaluated by measuring the ratio of the number of peaks overlapping with CTCF-FE and CTCF+FE (R_CTCF_). For IHO1 and HORMAD1, in all genotypes tested, this ratio was between 4.4 and 6.1, thus similar to the ratio of CTCF-FE to CTCF+FE (28134/4542=6.2) indicating that the peak number at CTCF+FE is mainly driven by CTCF enrichment. In contrast, SYCP3 peaks show an elevated R_CTCF_ (5.5 and 4.5) only in condition of DSB activity at hotspots (WT and *Hormad1KO*), but not in *Mei4KO*, *Spo11KO* and *Iho1KO* (Table S2). This supports the interpretation for increased occupancy of SYCP3 at FE in the absence of DSB activity and for a change of association of SYCP3 to FE and CTCF upon DSB formation.

### Determination of functional elements (FE)

First, promoters and enhancers, as defined in the Ensembl database, were merged. Of these sites, recombination hotspots (as defined by DMC1 SSDS-ChIP in WT(B6)) were excluded. Then, using the zygotene dataset of ^86^, H3K4me3 enrichment was scored within 10bp bins at +/-500bp from the center of FE. A FE was considered informative, when the mean enrichment score was ≥2. Last, the window around a given FE was extended to +/-1kb. This yielded 69963 peaks. We filtered the overlap with CTCF and obtained a pool of 57431 FE-CTCF peaks which was used for all quantifications on FE unless otherwise noted.

### Signal normalization and quantitative analysis

For most samples (see Table S1), biological duplicates were generated, analysed and read distributions and signal intensities were calculated after pooling reads from both replicates, if the IDR conditions were satisfied. If not, only one replicate was considered, (see above). Reads were normalized by library size in fragments per million (FPM) or reads per million (RPM) and with the corresponding knockout, except for CTCF ChIP data where the input was used. Read coverage was assessed with deeptools (version 3.4.1). Readcount was performed with the bedtools suite (version 2.26.0) and the intersect module.

### Selected intervals for browser images

Hotspots: Chr1; 13,840,000-13,880,000 and 168,140,000-168,180,000

CTCF sites: Chr1; 171,100,000-171,120,000 and 181,880,000-181,900,000

Functional elements: Chr1; 90,600,000-90,620,000 and 93,480,000-93,520,000 For all browsers, the Y axis is in RPM.

### Enrichment in regions with high copy number of the mo-2 minisatellite

The locations of mo-2 at the ends of several mouse chromosomes were mapped by Acquaviva et al. ^32^ who identified multiples copies in tandem at the ends of chr 4, 9, 13, X and Y. By Blast analysis, we also found two internal positions with more than two copies of mo-2 on chr 7 and 3. We thus defined 13 genomic positions with mo-2 (fig. S1C) that we used to quantify the enrichment of MEI4, IHO1, HORMAD1 and SYCP3. We did not include chr 4 as the genome assembly shows a large gap in the region of interest. The enrichment of MEI4, IHO1, HORMAD1 and SYCP3 in these regions in WT is quantified in fig. S1D and a browser window on chr X in fig. S1E. Several regions had signal close to background and were not included in WT vs mutant analysis (chr 3; X-1; X-2; Y-1). Region in chr 7 overlapped with a PRDM9 and DMC1 hotspot and was also not included this analysis. The enrichment of MEI4, IHO1, HORMAD1 and SYCP3 was quantified in mutant backgrounds showing that IHO1 enrichment depends on MEI4 at all sites tested whereas MEI4, HORMAD1 and SYCP3 were similarly enriched in WT and mutants (fig. S1F and G).

### Heatmaps

The heatmaps show the number of RPM, normalized to the ChIP signal in synchronized testes from ChIP samples of the respective KO mouse strains except for CTCF ChIP, which was normalized to the input. The enrichment was calculated in a −5 kb to +5 kb window around hotspot, CTCF or FE centres and averaged within 10bp bins. For all heatmaps, sites were ranked by decreasing intensity for the protein used to define these sites from top to bottom. The averaged profiles represent the normalized mean signal.

### Boxplots

For boxplot analyses, ChIP-seq signal intensity was calculated from pooled replicates (when the IDR conditions were satisfied), and normalized to the library size in FPM and to the corresponding KO mouse strain (i.e. *Mei4KO*, *Iho1KO*, *Hormad1KO* and *Sycp3KO*). For hotspots, the signal was measured at the 2000 strongest DMC1 sites within 1kb up– and down-stream of the hotspot centres (as defined by DMC1-SSDS) for MEI4, and within 700bp up and down-stream of the hotspot centers for IHO1, HORMAD1 and SYCP3. For CTCF sites, the signal was measured at the 5000 strongest CTCF sites (as defined by our CTCF ChIP-seq experiments) within 1kb up– and down-stream of the CTCF sites. For FE, the signal was measured at the 5000 strongest FE sites (as defined above), 1kb up or down-stream of the FE centres. All boxplots show the median and the 25^th^ to 75^th^ percentiles.

### Correlation plots

For hotspots, the peak intensity of all DMC1 sites ^31^ was correlated to the enrichment of MEI4 at 1kb up– and down-stream of hotspots and the enrichment of IHO1, HORMAD1 and SYCP3 at 2.5kb up– and down-stream of hotspots. For CTCF sites, the peak intensity of all CTCF sites without FE (as defined by our CTCF ChIP-seq analysis of synchronized testes at day 8 post-injection) were correlated to the enrichment of IHO1, HORMAD1 and SYCP3 within 1kb up– and down-stream of CTCF sites. For FE, the read enrichment of H3K4me3 of FE sites without CTCF was correlated to the read enrichment of IHO1, HORMAD1 and SYCP3 within 1kb up– and down-stream of FE. Log scales were used and values ≤0 were excluded. Correlations were evaluated with Rho, the Spearman correlation coefficient.

### Statistical analysis

Cytological data were analysed with GrapPad Prism 9. Significant differences in MEI4 focus count and intensity were assessed with the two-tailed Mann-Whitney test. P-values for significant differences between genotypes are indicated on the figures. Two or three mice per genotype were used. The number of nuclei per stage is indicated in the source data. For ChIP analysis, correlations were assessed with the two-tailed non-parametric Spearman correlation coefficient. Due to the large number of data points, non-parametric rank tests did not seem useful for comparative analyses between genotypes. Comparisons were performed by quantifying the mean difference relative to the standard deviation (SD) of the samples using the Cohen’s D parameter (D= |MeanA-MeanB|/SD(A,B). This parameter expresses differences in units of variability ^98^. Here, wild type (sample A) and mutant (sample B) samples were compared. When D is ≤0.2, the effect of the mutation is considered to be small. These quantifications are reported in Table S4.

## Supplemental Figures and Items

Figures S1-S7

Tables S1-S5

**Figure S1 related to Figure 1.**
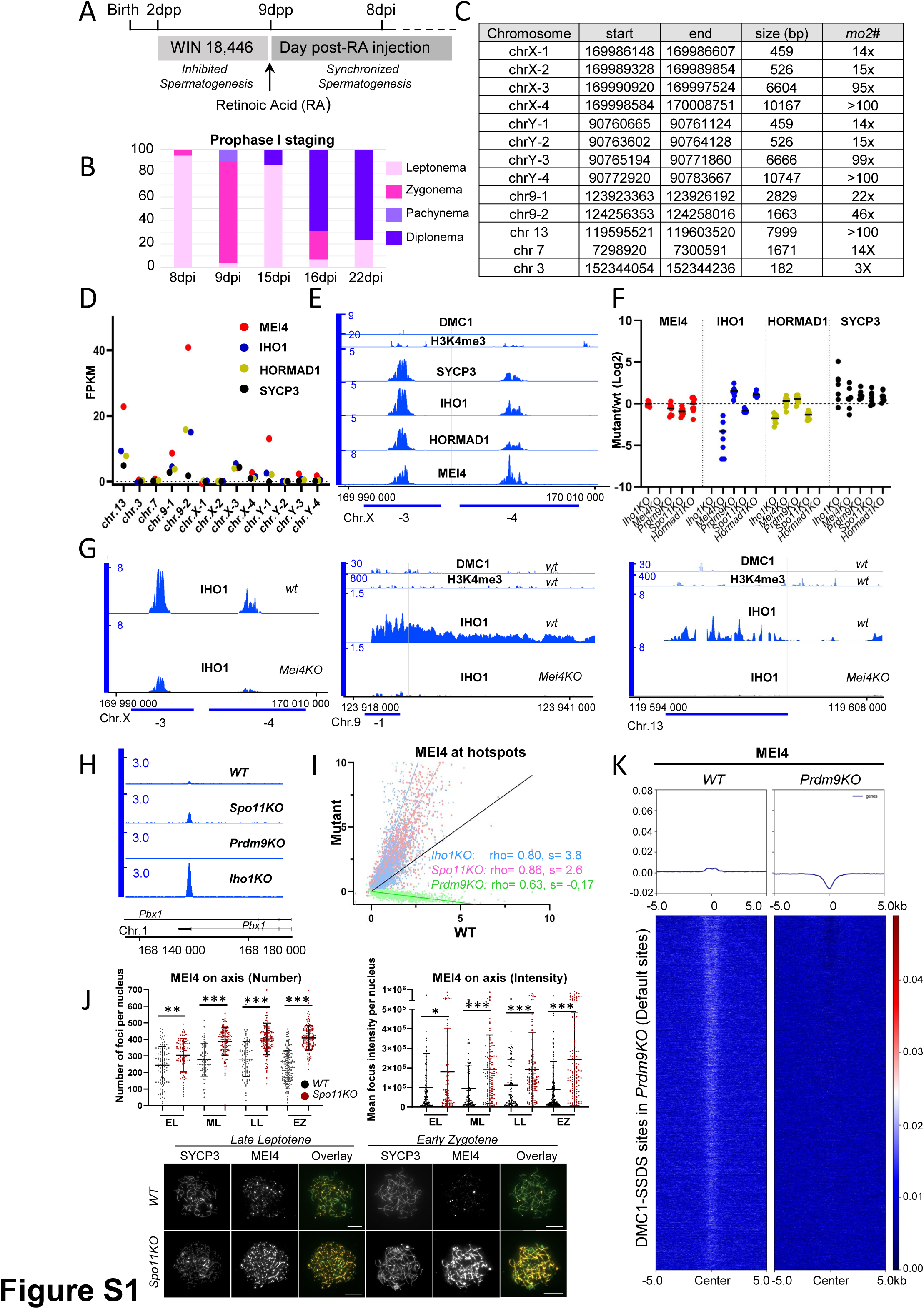
MEI4 is strongly recruited at *mo-2* repeat-containing regions and MEI4 enrichment at hotspots is SPO11– and IHO1-independent, but PRDM9-dependent. (A) Schematic representation of the prophase I spermatocyte synchronization protocol used for all ChIP experiments and in all mouse strains. (B) Cytological spermatocyte staging in wild type (*B6)* mice, showing the percentage of nuclei (SYCP3 positive) at each stage of prophase I at different days after retinoic acid injection (day post-injection, dpi). The staging criteria are described in the Star Methods section. (C) Summary table indicating the genomic position and number of *mo2* repeats in chromosomes (chr) X, Y, 9, 13, 7 and 3. (D) Enrichment of MEI4 (red), IHO1 (blue), HORMAD1 (green) and SYCP3 (black) within *mo2-*containing regions, normalized to the fragment length and library size. (E) ChIP-seq read distribution of DMC1-SSDS ^31^ and H3K4me3 ^86^, SYCP3, IHO1, HORMAD1 and MEI4 in wild type spermatocytes within two *mo2-*containing regions (chrX-3, chrX-4) on chromosome X. (F) Enrichment of MEI4, IHO1, HORMAD1 and SYCP3 at *mo2* sites in *Iho1KO*, *Mei4KO*, *Prdm9KO*, *Spo11KO* and *Hormad1KO* spermatocytes relative to the wild type sample. Enrichment was assessed in a pool of 8 *mo2*-containing regions (see Star Methods). (G) Left: ChIP-seq read distribution of IHO1 in wild type (*wt*) and *MEI4KO* spermatocytes in two *mo2-*containing regions (chrX-3, chrX-4) on chromosome X. Middle: ChIP-seq read distribution of DMC1-SSDS ^31^, H3K4me3 ^86^, and IHO1 in *wt* and *Mei4KO* within a *mo2*-containing region on chromosome 9. Right: ChIP-seq read distribution of DMC1-SSDS, H3K4me3 and IHO1 in *wt* and *Mei4KO* spermatocytes in a *mo2*-containing region on chromosome 13. (H) MEI4 read distribution at a representative hotspot in wild type (WT) and in *Spo11KO*, *Prdm9KO* and *Iho1KO* spermatocytes. (I) MEI4 signal intensity (FPM) at *Dom2* hotspots in *Iho1KO*, *Spo11KO* and *Prdm9KO* compared to WT. Rho, Spearman correlation coefficient; s, slope of the linear regression. (J) Top left: Number of MEI4 foci in early leptonema (EL), mid leptonema (ML), late leptonema (LL), and early zygonema (EZ) or zygonema-like stages in *WT* (black) *vs Spo11KO* (pink) synchronized spermatocytes. Each dot represents the number of MEI4 foci per nucleus (mean ± SD). Top right: Quantification of mean MEI4 focus intensity per nucleus (mean ± SD; a.u., arbitrary units) in *WT* (black) vs *Spo11KO* (pink) mice at same stages as above. P values were assessed by Man-Whitney test (* p = 0.0017, ** p= 0.002, *** p< 0.0001). Bottom: representative images of MEI4 staining at late leptonema and early zygonema in wild type vs *Spo11KO* spermatocyte spread nuclei at 8dpi (same animals as for ChIP), scale bar indicates 10µm. (K) MEI4 ChIP-seq signal in *WT* and *Prdm9KO* mice at DMC1-SSDS sites in *Prdm9KO* (default sites) ^99^.

**Figure S2 related to Figure 1.**
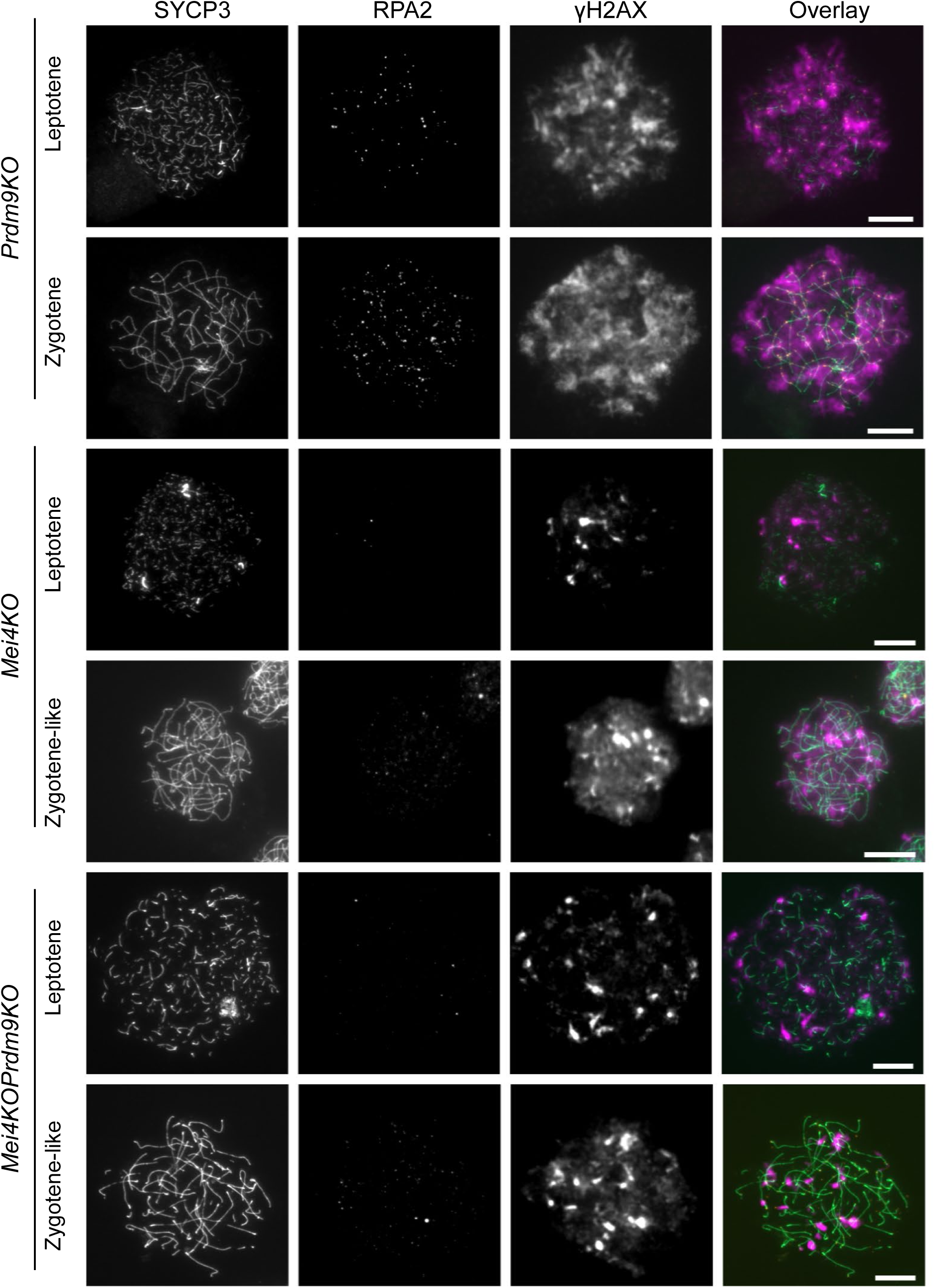
MEI4 is essential for DSB formation in *Prdm9KO*. Immunofluorescence staining for SYCP3, RPA and γH2AX on spread spermatocytes from testes of adult mice, showing representative nuclei of leptotene, zygotene or zygotene-like nuclei in *Prdm9KO* (two upper rows), *Mei4KO* (two middle rows), and *Mei4KO Prdm9KO* (two lower rows) mice.

**Figure S3 related to Figure 1.**
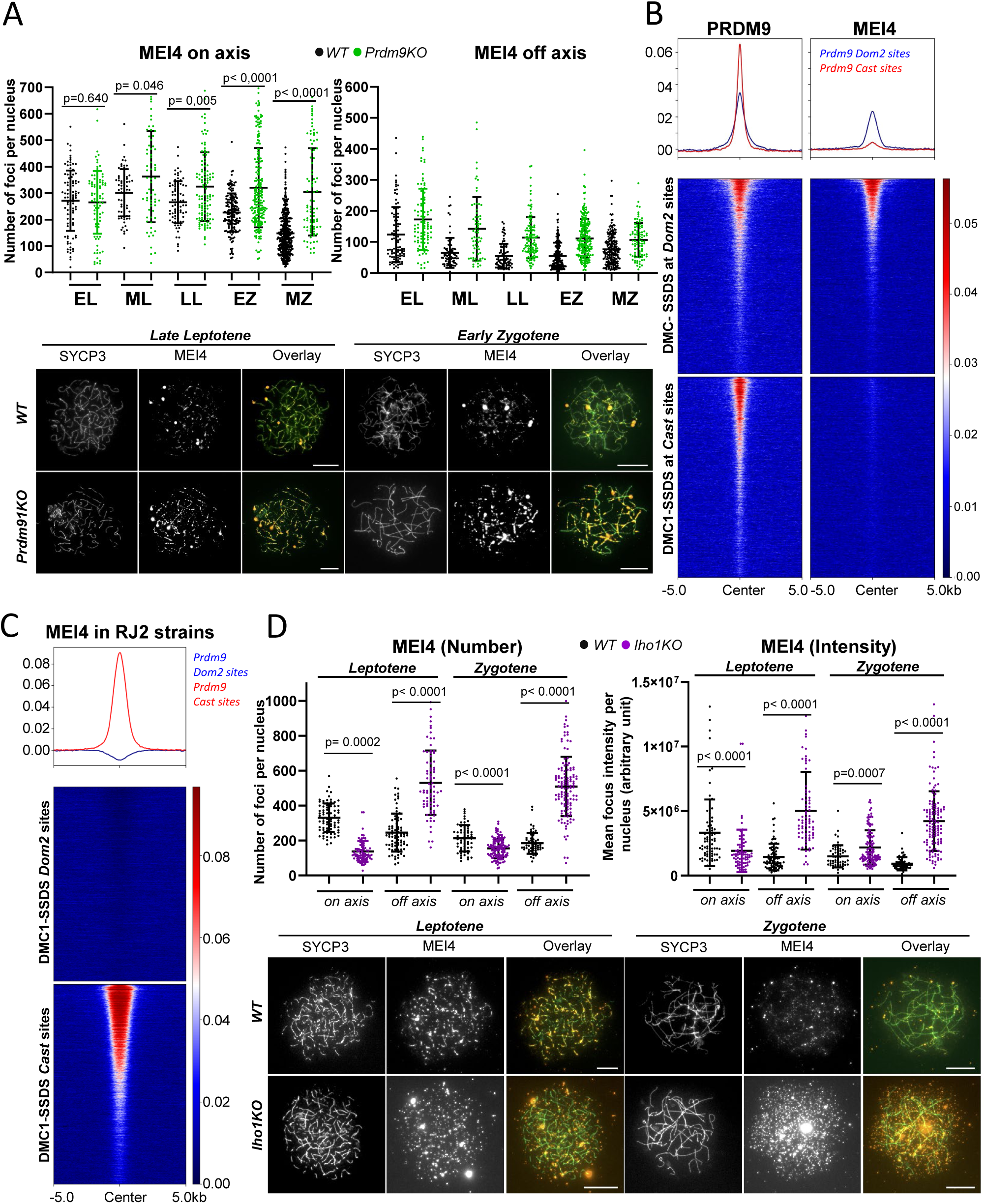
MEI4 recruitment at hotspots and axes is dependent on PRDM9 catalytic activity and IHO1, respectively. (A) MEI4 focus formation in wild type (WT) (black) and *Prdm9KO* (green) spermatocyte nuclei. Top left: Number of MEI4 foci that co-localize on SYCP3-positive axes in early leptonema (EL), mid leptonema (ML), late leptonema (LL), early zygonema (EZ), and mid zygonema (MZ). Top right: Number of MEI4 foci outside SYCP3-positive axes. Each dot represents the number of MEI4 foci per nucleus (mean ± SD). Bottom: representative images of MEI4 staining at late leptonema and early zygonema in wild type vs *Prdm9KO* spermatocyte spread nuclei at 8dpi (same animals as for ChIP), scale bar indicates 10µm. (B) PRDM9 (left panels) and MEI4 (right panels) ChIP-seq signals in *B6*-Tg(YF) at *Dom2* (top) and *Cast* (bottom) sites, defined by DMC1-SSDS in *B6* and *RJ2* mice, respectively^31^. (C) MEI4 ChIP-seq signal from RJ2 mice, expressing the *Prdm9^Cst^* allele, at *Dom2* (top) and *Cast* (bottom) sites defined by DMC1-SSDS in *B6* and *RJ2* mice ^31^. The dip at the centre of the *Dom2* average enrichment profile is due to a stronger background of the ChIP-seq signal in *Mei4KO* (expressing PRDM9*^Dom^*^2^) mice used for normalization. (D) MEI4 focus formation in WT (black) and *Iho1KO* (purple) spread spermatocyte nuclei from day 12dpp testes. Top left: number of MEI4 foci in leptonema and early zygonema or zygonema-like. Number of foci that colocalize with SYCP3-positive axes (on-axis) and that are outside SYCP3-positive axes (off-axis). Top right: Quantification of mean MEI4 focus intensity (mean ± SD; a.u., arbitrary units) in WT (black) *vs Iho1KO* (purple) mice at the same stages as in the left panel. Signal intensity for foci that localize on SYCP3-positive axes (on-axis) and for off-axis foci. Bottom: representative images of MEI4 staining in leptonema and zygonema in wild type vs *Iho1KO* 14dpp spermatocyte spread nuclei, scale bar indicates 10µm.

**Figure S4 related to Figure 2.**
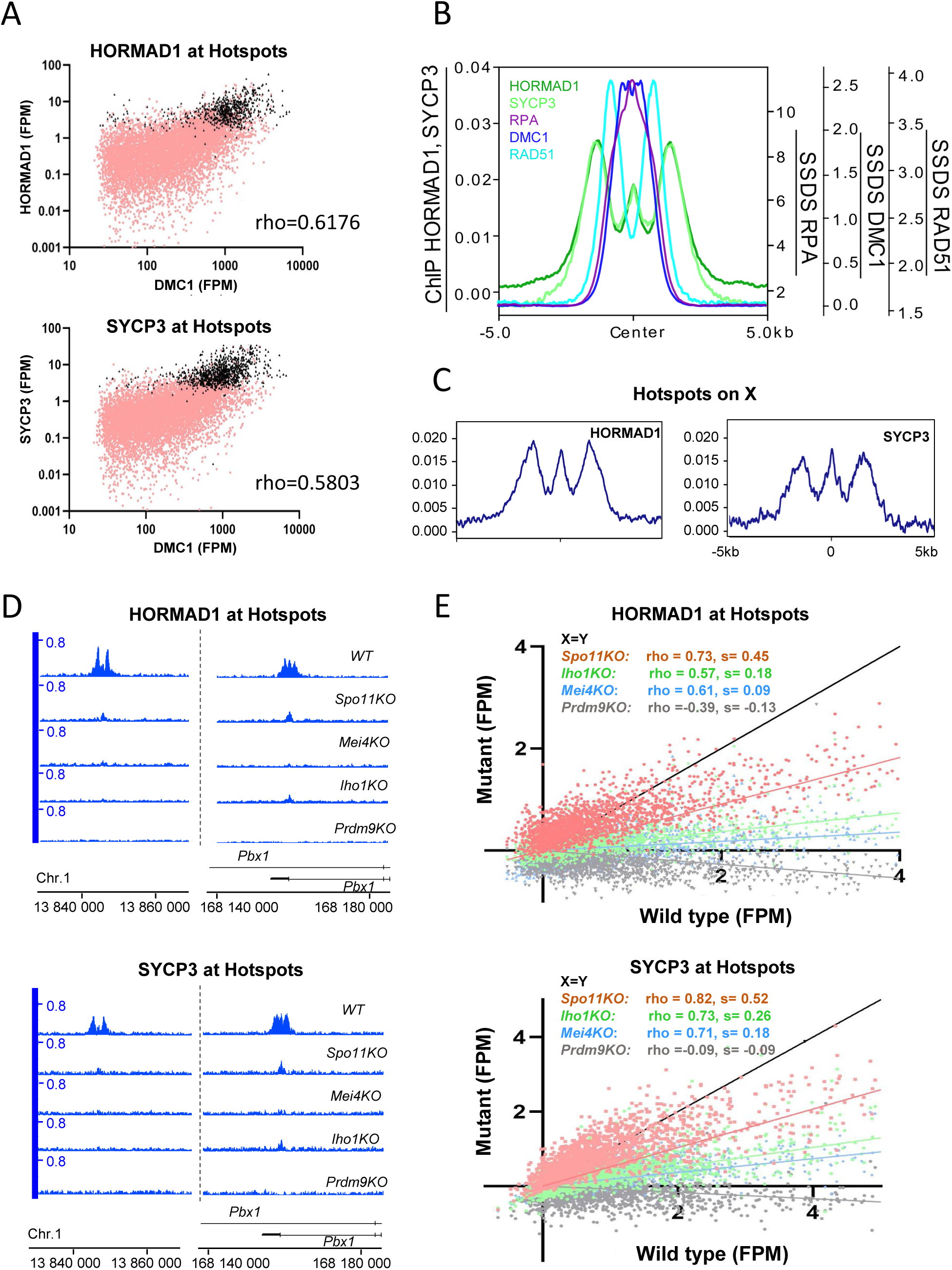
HORMAD1 and SYCP3 recruitment at hotspots is DSB-independent, but PRDM9, – MEI4– and IHO1-dependent. (A) HORMAD1(top) and SYCP3 (bottom) signals compared with DMC1-SSDS signal at all *Dom2* DSB hotspots in B6 mice ^31^. Black and pink dots highlight peaks that overlap and that do not overlap with hotspots, respectively. Rho: Spearman correlation coefficient. (B) Averaged and normalized enrichment profile of HORMAD1 and SYCP3 ChIP-seq, compared to the averaged enrichment profile of RPA, DMC1 and RAD51 recombinase at *Dom2* hotspots ^100^. (C) Averaged and normalized enrichment profile of HORMAD1 and SYCP3 ChIP-seq at non-PAR hotspots on chromosome X (D) ChIP-seq read distribution of HORMAD1 (top) and SYCP3 (bottom) in wild type (WT), *Spo11KO*, *Mei4KO*, *Iho1KO* and *Prdm9KO* spermatocytes at two representative hotspots. (E) HORMAD1 and SYCP3 signal intensity (FPM) at *Dom2* hotspots in *Iho1KO*, *Spo11KO*, *Mei4KO*, *Iho1KO* and *Prdm9KO* compared with WT spermatocytes. Rho: Spearman correlation coefficient. s: slope of the linear regression.

**Figure S5 related to Figure 3.**
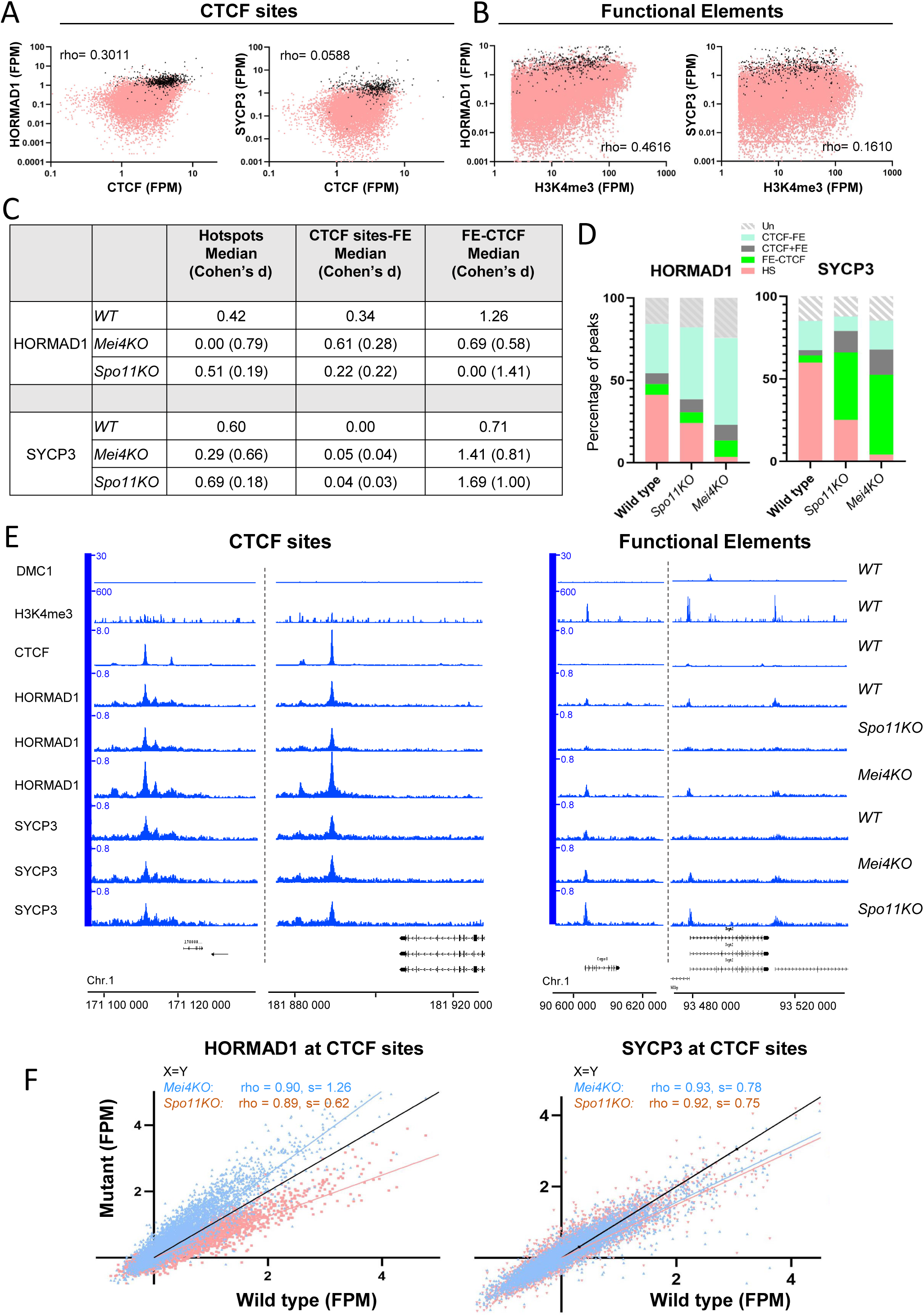
Distinct HORMAD1 and SYCP3 binding dynamics at CTCF sites and functional elements (FE). (A) HORMAD1 and SYCP3 signals (FPM) compared with the CTCF signal intensity (FPM) at CTCF sites in wild type (WT) spermatocytes. Black and pink dots highlight HORMAD1 or SYCP3 peaks that overlap and that do not overlap with CTCF peaks, respectively. Rho: Spearman correlation coefficient. (B) HORMAD1 and SYCP3 signals (FPM) compared with the H3K4me3 signal intensity ^86^ at FE in WT spermatocytes. Black and pink dots highlight HORMAD1 or SYCP3 peaks that overlap and that do not overlap with FE, respectively. Rho: Spearman correlation coefficient. (C) Median HORMAD1 and SYCP3 signal intensity in WT, *Mei4KO* and *Spo11KO* at hotspots (2000 sites), CTCF sites (CTCF-FE, 5000 sites) and FE (FE-CTCF, 5000 sites). Quantitative differences were assessed with the Cohen’s coefficient “d” (see Star Methods). Differences were considered as small when d ≤ 0.2. (D) Percentage of peaks in the wild type, *Spo11KO* and *Mei4KO* strains at five different genomic site types: hotspots (HS), FE-CTCF, CTCF+FE, CTCF-FE, and undefined sites (Un). (E) ChIP-seq read distribution of DMC-SSDS, H3K4me3 ^86^, CTCF, HORMAD1 and SYCP3 in wild type (WT), compared to HORMAD1 and SYCP3 read distribution in the *Spo11KO* and *Mei4KO* strains at two representative CTCF sites and three representative functional elements. (F) HORMAD1 and SYCP3 signal intensity (FPM) at CTCF sites in the *Mei4KO* and *Spo11KO* strains compared with wild type. Rho: Spearman correlation coefficient. s: slope of the linear regression.

**Figure S6 related to Figure 4.**
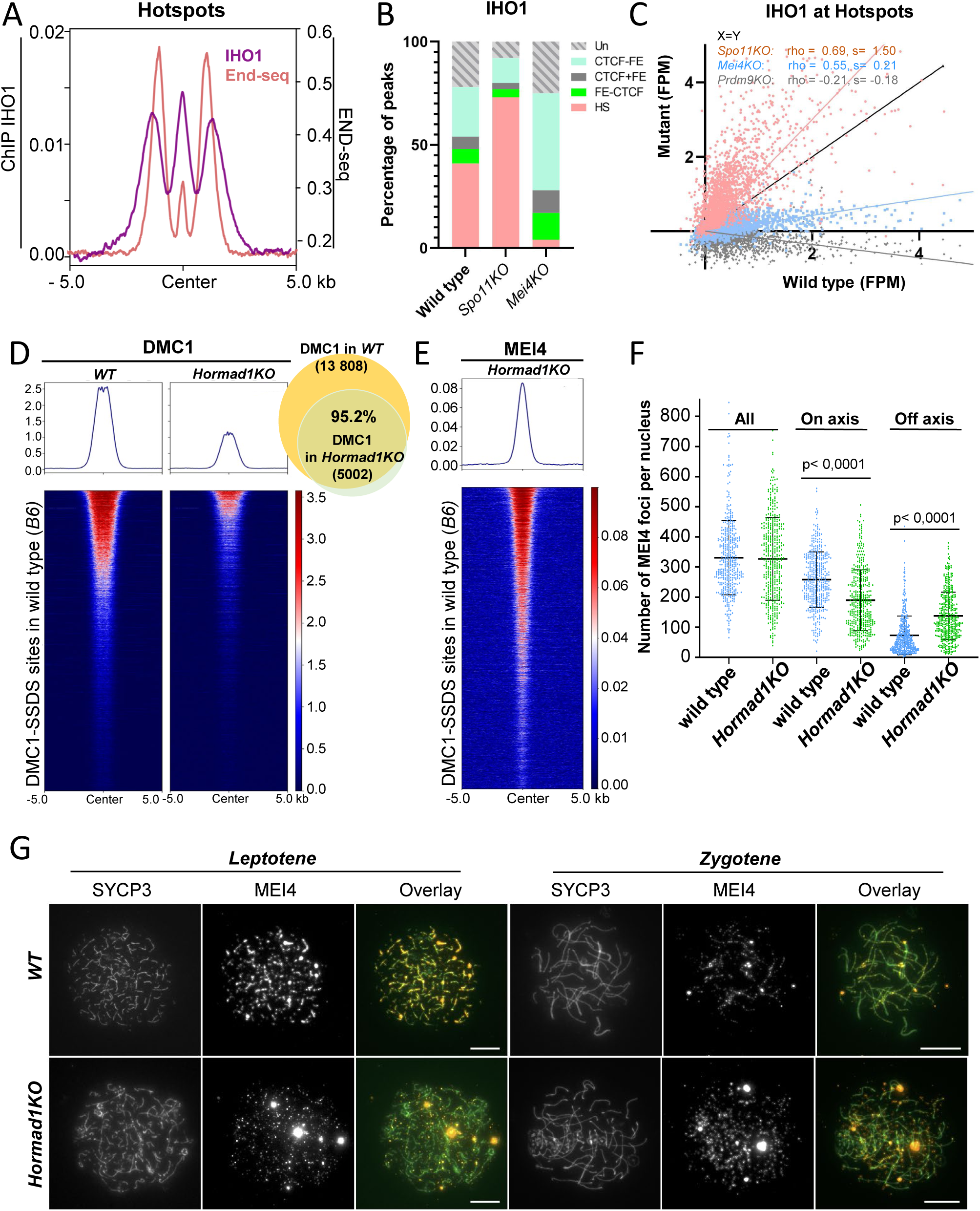
IHO1, a link between DSB proteins and axis proteins at hotspots. (A) Averaged enrichment of IHO1 (ChIP-seq) (purple) and resection track ends, assessed by END-seq (red) ^48^, at *Dom2* hotspots. (B) Percentage of peaks in the wild type, *Spo11KO* and *Mei4KO* strains at five different genomic site types: hotspots (HS), FE-CTCF, CTCF+FE, CTCF-FE, and undefined sites (Un). (C) IHO1 signal intensity (FPM) at *Dom2* hotspots ^31^ in the *Spo11KO*, *Mei4KO* and *Prdm9KO* strains compared with wild type. Rho: Spearman correlation coefficient. s: slope of the linear regression. (D) Left: DMC1-SSDS signal in wild type and *Hormad1KO* synchronized testes at *Dom2* hotspots ^31^. Right: Venn diagram showing the overlap of DMC1-SSDS peaks in WT (yellow) and DMC1-SSDS peaks in *Hormad1KO* mice (light green). (E) MEI4 ChIP-seq signal in *Hormad1KO* spermatocytes at *Dom2* hotspots. (F) MEI4 focus formation in wild type and *Hormad1KO* spread spermatocyte nuclei from 8dpi testes. MEI4 foci were assessed from early leptonema to early zygonema (pooled). Left: total number of foci (All). Middle: number of foci that colocalize with SYCP3-positive axes (on-axis). Right: number of foci that localize outside SYCP-positive axes (off-axis). Each dot represents the number of MEI4 foci per nucleus (mean ± SD). (G) Representative images of MEI4 staining at leptonema and zygonema in wild type vs *Hormad1KO* spermatocyte spread nuclei at 8dpi (same animals as for ChIP), scale bar indicates 10µm.

**Figure S7 related to Figure 5.**
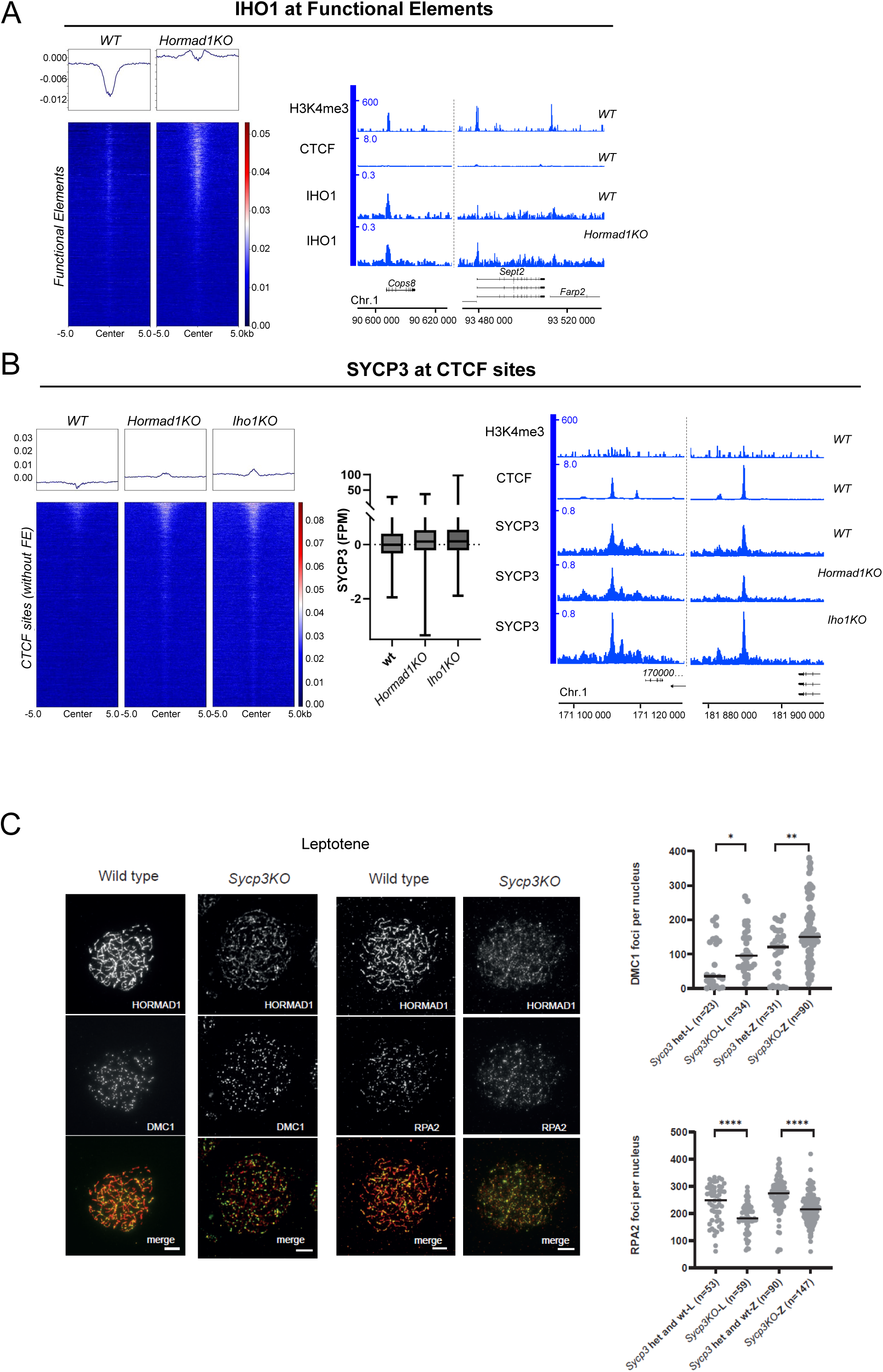
SYCP3 and IHO1 bind weakly to CTCF sites and FE respectively. (A) Left: IHO1 ChIP-seq signal at FE in WT and *Hormad1KO*. Right: ChIP-seq read distribution of IHO1, H3K4me3 ^86^ and CTCF in the WT and *Hormad1KO* mouse strains at three representative FE. (B) Left: SYCP3 ChIP-seq signal at CTCF sites in the wild type (WT), *Hormad1KO* and *Iho1KO* strain. Middle: SYCP3 ChIP-seq signal intensity at CTCF sites in WT, *Hormad1KO*, and *Iho1KO* spermatocytes. The signal was measured at the 5000 strongest CTCF sites. Right: ChIP-seq read distribution of SYCP3, H3K4me3 ^86^, and CTCF in the WT, *Hormad1KO* and *Iho1KO* mouse strains at two representative CTCF sites. (C) Immunolocalisation of HORMAD1, DMC1 and RPA2 in wild type and *Sycp3KO* spermatocytes at leptonema. The scale bar indicates 10µm (left). Quantification of the number of foci per nucleus in wild type and *Sycp3KO* spermatocytes at leptonema (right). P values were assessed by a Mann-Whitney test (*p < 0.05, **p < 0.01, ***p < 0.001, ****p < 0.0001).

**Supplemental Table S1.** related to Methods. List of all ChIP-seq experiments used in this study. Prophase I staging was performed as described in Materials and Methods. The percentage of prophase stages among SYCP3 positive cells are indicated. L indicates leptonema and Z indicates zygonema. Star (*) indicates samples where only one replicate was analysed, because of experimental reasons or because the other replicate did not pass our reproducibility criteria (see Methods).

| Protein (ChIP) | Mouse strain | Prophase I Staging | Sample Name in GEO |
| --- | --- | --- | --- |
| PRDM9 | wild type (B6) | 3%L 97%Z | B_WT_P9_r1 |
| PRDM9 | wild type (B6) | 13.7%L 86.3%Z | B_WT_P9_r2 |
| PRDM9 | Prdm9KO | 44.4%L 55.6%Z | P9_KO_P9_r1 |
| PRDM9 | Prdm9KO | 86.9%L 13.1%Z | P9_KO_P9_r2 |
| PRDM9* | B6-Tg(YF) | 16.3%L 83.7%Z | Yf_Tg_P9 |
| MEI4 | wild type (B6) | 47.5%L 52.5%Z | B_WT_M4_r1 |
| MEI4 | wild type (B6) | 47.5%L 52.5%Z | B_WT_M4_r2 |
| MEI4 | Prdm9KO | 31.3%L 68.7%Z | P9_KO_M4_r1 |
| MEI4 | Prdm9KO | 44.4%L 55.6%Z | P9_KO_M4_r2 |
| MEI4 | B6-Tg(YF) | 16.3%L 83.7%Z | YF_Tg_M4_r1 |
| MEI4 | B6-Tg(YF) | 50%L 50%Z | YF_Tg_M4_r2 |
| MEI4* | RJ2 | 58.6%L 41.4%Z | RJ_M4 |
| MEI4 | Spo11KO | 44%L 56%Z | S11_KO_M4_r1 |
| MEI4 | Spo11KO | 73.7%L 26.3%Z | S11_KO_M4_r2 |
| MEI4 | Mei4KO | 100% L | M4_KO_M4_r1 |
| MEI4 | Mei4KO | 97.1% L 2.9%Z | M4_KO_M4_r2 |
| MEI4 | Iho1KO | 70%L 30%Z | I1_KO_M4_r1 |
| MEI4 | Iho1KO | 74.2%L 25.8%Z | I1_KO_M4_r2 |
| MEI4 | Hormad1KO | 53.7%L 46.3%Z | H1_KO_M4_r1 |
| MEI4 | Hormad1KO | 9%L 91%Z | H1_KO_M4_r2 |
| IHO1 | wild type (B6) | 3%L 97%Z | B_WT_I1_r1 |
| IHO1 | wild type (B6) | 13.7%L 86.3%Z | B_WT_I1_r2 |
| IHO1 | Prdm9KO | 86.9%L 13.1%Z | P9_KO_I1_r1 |
| IHO1 | Prdm9KO | 14.8%L 85.2%Z | P9_KO_I1_r2 |
| IHO1 | Spo11KO | 73.7%L 26.3%Z | S11_KO_I1_r1 |
| IHO1 | Spo11KO | 55.5%L 44.5%Z | S11_KO_I1_r2 |
| IHO1 | Mei4KO | 75%L 25%Z | M4_KO_I1_r1 |
| IHO1 | Mei4KO | 70.4%L 29.6%Z | M4_KO_I1_r1 |
| IHO1 | Hormad1KO | 9%L 91%Z | H1_KO_I1_r1 |
| IHO1 | Hormad1KO | 46.2%L 53.8%Z | H1_KO_I1_r2 |
| IHO1 | Iho1KO | 70%L 30%Z | I1_KO_I1_r1 |
| IHO1 | Iho1KO | 74.2%L 25.8%Z | I1_KO_I1_r2 |
| HORMAD1 | wild type (B6) | 100%Z | B_WT_H1_r1 |
| HORMAD1 | wild type (B6) | 3%L 97%Z | B_WT_H1_r2 |
| HORMAD1 | Prdm9KO | 31.3%L 68.7%Z | P9_KO_H1_r1 |
| HORMAD1 | Prdm9KO | 44.4%L 55.6%Z | P9_KO_H1_r2 |
| HORMAD1 | Spo11KO | 73.7%L 26.3%Z | S11_KO_H1_r1 |
| HORMAD1 | Spo11KO | 100%L | S11_KO_H1_r2 |
| HORMAD1 | Mei4KO | 100% L | M4_KO_H1_r1 |
| HORMAD1 | Mei4KO | 97.1% L 2.9%Z | M4_KO_H1_r2 |
| HORMAD1 | Iho1KO | 70%L 30%Z | I1_KO_H1_r1 |
| HORMAD1 | Iho1KO | 74.2%L 25.8%Z | I1_KO_H1_r2 |
| HORMAD1 | Hormad1KO | 53.7%L 46.3%Z | H1_KO_H1_r1 |
| HORMAD1 | Hormad1KO | 9%L 91%Z | H1_KO_H1_r2 |
| SYCP3 | wild type (B6) | 13.7%L 86.3%Z | B_WT_S3_r1 |
| SYCP3 | wild type (B6) | 5.3%L 94.7%Z | B_WT_S3_r2 |
| SYCP3* | Prdm9KO | 86.9%L 13.1%Z | P9_KO_S3 |
| SYCP3* | Spo11KO | 55.5%L 44.5%Z | S11_KO_S3 |
| SYCP3 | Mei4KO | 70.4%L 29.6%Z | M4_KO_S3_r1 |
| SYCP3 | Mei4KO | 100%L | M4_KO_S3_r2 |
| SYCP3* | Iho1KO | 74.2%L 25.8%Z | I1_KO_S3 |
| SYCP3 | Hormad1KO | 46.2%L 53.8%Z | H1_KO_S3_r1 |
| SYCP3 | Hormad1KO | 23.6%L 76.4%Z | H1_KO_S3_r2 |
| SYCP3* | Sycp3KO | 43%L 57%Z | S3_KO_S3 |
| CTCF | wild type (B6) | 5.3%L 94.7%Z | B_WT_CTCF_r1 |
| CTCF | wild type (B6) | 22.1%L 77.9%Z | B_WT_CTCF_r2 |
| CTCF | Prdm9KO | 14.8%L 85.2%Z | P9_KO_CTCF_r1 |
| CTCF | Prdm9KO | 14.8%L 85.2%Z | P9_KO_CTCF_r2 |
| CTCF | Spo11KO | 100%L | S11_KO_CTCF_r1 |
| CTCF | Spo11KO | 100%L | S11_KO_CTCF_r2 |
| DMC1 SSDS | wild type (B6) | 17% L, 68% Z | B_WT_DMC1_r1 |
| DMC1 SSDS | wild type (B6) | 58% L, 42% Z | B_WT_DMC1_r2 |
| DMC1 SSDS | Hormad1KO | 51% L, 49% Z | H1_KO_DMC1_r1 |
| DMC1 SSDS | Hormad1KO | 50%L, 50%Z | H1_KO_DMC1_r2 |
| INPUT* | wild type (B6) | 22.1%L 77.9%Z | Inp_B6 |

**Supplemental Table S2.** related to Figures 1-4. **Number of retrieved peaks.** Summary of the number of retrieved peaks in MEI4, IHO1, HORMAD1 and SYCP3 ChIP experiments in wild type (*B6*) and mutant synchronized testes samples within the different types of sites (hotspots, CTCF sites, CTCF sites coinciding with functional elements (FE), FE and undefined sites.

| Protein | Peaks | *B6 (WT)* | *Prdm9KO* | *Mei4KO* | *Iho1KO* | *Spo11KO* | *Hormad1KO* |
| --- | --- | --- | --- | --- | --- | --- | --- |
| **MEI4** | **Total** | 1566 | 80 | n.a | 2679 | 3817 | 3828 |
|  | Hotspots | 1511 | 6 | n.a | 2626 | 3772 | 3719 |
|  | FE-CTCF | 8 | 10 | n.a | 8 | 6 | 23 |
|  | CTCF+FE | 2 | 3 | n.a | 4 | 2 | 10 |
|  | CTCF-FE | 13 | 18 | n.a | 11 | 11 | 16 |
|  | Unidentified | 32 | 43 | n.a | 30 | 26 | 60 |
| **IHO1** | **Total** | 1589 | 378 | 1036 | na | 2681 | 1860 |
|  | Hotspots | 647 | 17 | 37 | n.a | 1954 | 1739 |
|  | FE-CTCF | 113 | 33 | 130 | n.a | 114 | 34 |
|  | CTCF+FE | 87 | 32 | 111 | n.a | 71 | 14 |
|  | CTCF-FE | 385 | 197 | 491 | n.a | 318 | 13 |
|  | Unidentified | 357 | 99 | 267 | n.a | 224 | 60 |
| **HORMAD1** | **Total** | 1715 | 1009 | 5275 | 3149 | 2897 | n.a |
|  | Hotspots | 706 | 26 | 191 | 171 | 698 | n.a |
|  | FE-CTCF | 112 | 109 | 524 | 227 | 187 | n.a |
|  | CTCF+FE | 115 | 118 | 500 | 299 | 230 | n.a |
|  | CTCF-FE | 511 | 527 | 2785 | 1709 | 1266 | n.a |
|  | Unidentified | 271 | 229 | 1275 | 743 | 516 | n.a |
| **SYCP3** | **Total** | 1911 | 577 | 1125 | 5380 | 2374 | 823 |
|  | Hotspots | 1144 | 34 | 47 | 207 | 598 | 181 |
|  | FE-CTCF | 83 | 225 | 543 | 3098 | 969 | 83 |
|  | CTCF+FE | 61 | 72 | 172 | 696 | 309 | 73 |
|  | CTCF-FE | 338 | 105 | 197 | 602 | 210 | 330 |
|  | Unidentified | 285 | 141 | 166 | 777 | 288 | 156 |

**Supplemental Table S3.**
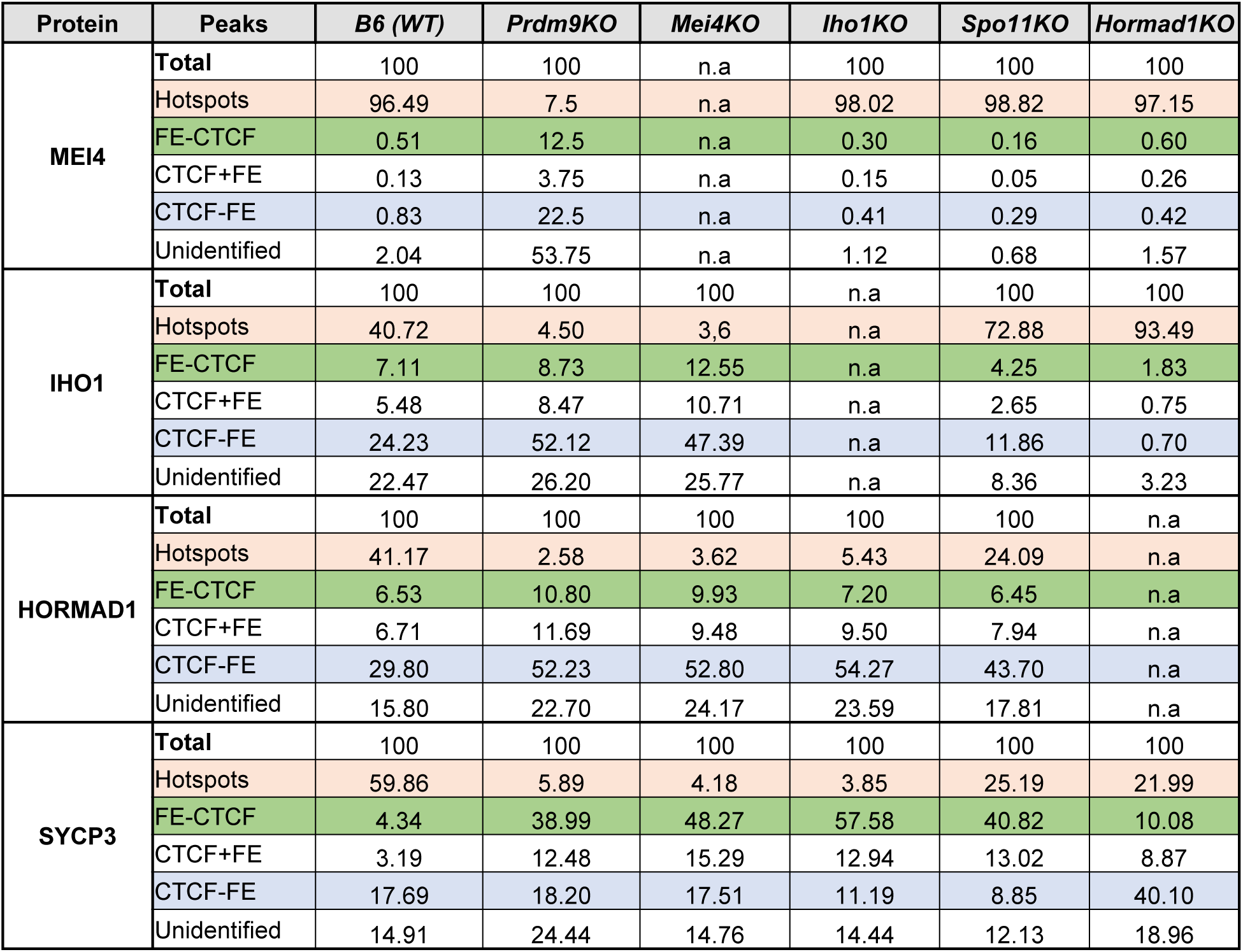
related to Figures 1-4. **Proportion of retrieved peaks.** Summary of de proportion of retrieved peaks for each type of site relative to the total number of peaks in MEI4, IHO1, HORMAD1 and SYCP3 ChIP experiments in wild type (*B6*) and mutant synchronized testes samples.

**Supplemental Table S4.**
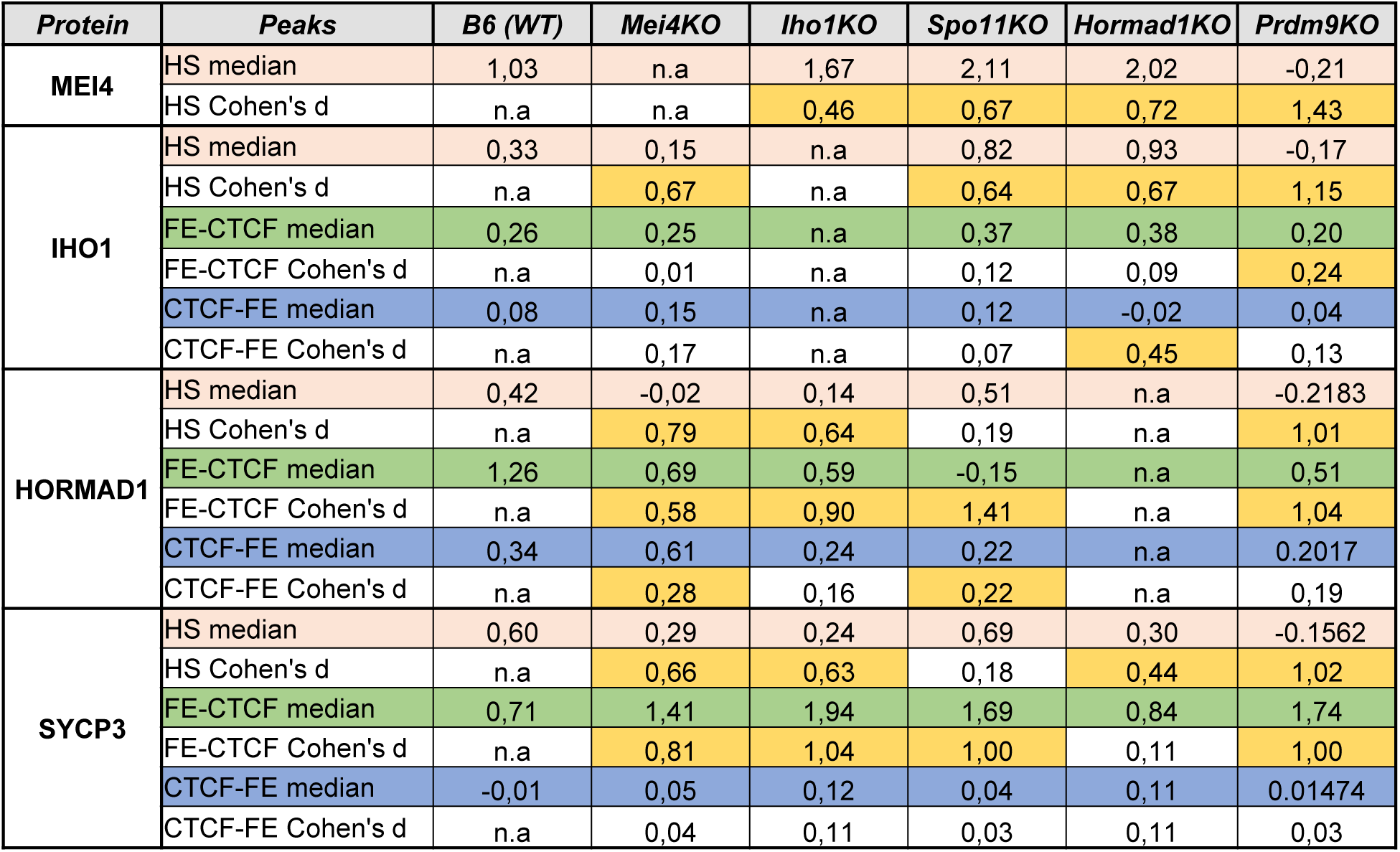
related to Figures 1E, 2E, 3E, 4D, 5. Median intensities and Quantitative differences. Summary of the median enrichment (FPM) of MEI4, IHO1, HORMAD1 and SYCP3 ChIP experiments computed over hotspots (2000 sites), CTCF sites (CTCF-FE, 5000 sites) and functional elements (FE-CTCF, 5000 sites). Quantitative differences were assessed by Cohen’s coefficient “d” (see STAR Methods). Differences were considered as important when d > 0.2 (shaded in yellow).

**Supplemental Table S5.** related to STAR Methods. Description of the mutant phenotypes of used mouse strains.

| Mouse strain | DSB formation | DSB localization | Axis formation | Synaptonemal complex formation | Stage of meiotic arrest | References |
| --- | --- | --- | --- | --- | --- | --- |
| *Prdm9KO* | Yes | Functional elements | Yes | Partial, homologous and non-homologous synapsis | Pseudo-pachytene | Hayashi et al. (2005) Nature 438, 374-378. |
| *Prdm9-YF-Tg* | Yes | Functional elements | Yes | Partial, homologous and non-homologous synapsis | Pseudo-pachytene | Diagouraga et al. (2018) Mol Cell 69, 853-865 e856. |
| *Mei4KO* | No | - | Yes | Partial, non-homologous synapsis | Pseudo-zygotene | Kumar et al. (2010) Genes Dev 24, 1266-1280. |
| *Iho1KO* | No | - | Yes | Partial, non-homologous synapsis | Pseudo-zygotene | Stanzione et al. (2016) Nat Cell Biol 18, 1208-1220. |
| *Spo11KO* | No | - | Yes | Partial, non-homologous synapsis | Pseudo-zygotene | Baudat et al. (2000) Mol Cell 6, 989-998.<br>Romanienko et al. (2000) Mol Cell 6(5):975-87. |
| *Hormad1KO* | 4 to 5 fold reduction | PRDM9-sites | Yes | Partial homologous synapsis | Pseudo-pachytene | Daniel et al. (2011) Nat Cell Biol 13, 599-610 |
| *Sycp3KO* | Yes | PRDM9-sites (predicted) | Cohesin core with discontinuous HORMAD1 | Short stretches of SYCP1 on cohesin cores, likely homologous | Pseudo-zygotene | Yuan et al. (2000) Molecular Cell 5, 73-83.<br>Peltari et al. (2001) MCB, 5667-5677.<br>This study |

